# Epithelial mesenchymal transition initiates precancer states in *BRCA1* mutation carriers

**DOI:** 10.64898/2026.05.06.723061

**Authors:** Neta Bar-Hai, Rakefet Ben-Yishay, Sheli Arbili-Yarhi, Rinat Bernstein-Molho, Gil Goldinger, Nora Balint-Lahat, Tehillah Menes, Naama Herman, Vered Noy, Aiham Mansour, Opher Globus, Pnina Hilman, Yonathan Zehavi, Inbal Eizenberg-Magar, Elmir Mahammadov, Thomas Conrad, Nikolaus Rajewsky, Yaron E. Antebi, Raanan Berger, Dana Ishay-Ronen

## Abstract

Epithelial-to-mesenchymal transition (EMT) is activated to equip cells with the capacity to adapt to and escape hostile conditions. While EMT is required for cancer progression, its role in breast cancer initiation remains elusive. Given the basal-like phenotype of breast cancers arising in female carriers of germline *BRCA1* pathogenic variants (*BRCA1* carriers), we hypothesized that enhanced EMT susceptibility underlies precancerous initiation in mammary epithelium. Perturbation of patient-derived normal mammary organoids from *BRCA1* carriers and non-carriers with inflammatory cytokines induced copy number variations (CNV) and the acquisition of oncogenic mutations in both groups. However, in organoids derived from *BRCA1* carriers, cytokine exposure induced morphological, transcriptomic, and functional EMT, accompanied by a transition to basal-like phenotype. Concomitant DNA damage accumulation in organoids from *BRCA1*-carriers demonstrated PARP inhibitor sensitivity. EMT-primed states were identified in a subpopulation of normal mammary epithelium from *BRCA1* carriers. We demonstrate the utility of patient-derived normal *BRCA1* heterozygous mammary organoids to reveal a plastic, high-risk epithelial state that is associated with a transient, targetable vulnerability.

## Main

Germline *BRCA1* pathogenic variants (PV) predispose female carriers to over 70% lifetime risk of developing aggressive triple-negative, basal like breast cancer^1,2^. *BRCA1*-associated breast cancer is characterized by BRCA1 dysfunction, resulting in deficient homologous recombination (HR) and accumulation of DNA damage^2,3^. While the impaired function of BRCA1 in breast cancer is broadly studied, it remains unclear what triggers genomic instability and epithelial plasticity in *BRCA1* heterozygous mammary epithelium^4–10^.

Modeling precancer events in a germline *BRCA1*heterozygous background is challenging^2,11^. Transgenic mouse models with heterozygous germline *BRCA1*-PV do not develop spontaneous tumors^11–13^. Therefore, *BRCA1* murine models require the somatic activation of an additional oncogenic mutation (typically in *TP53*) for tumorigenesis initiation^9,12^. Thus, processes leading to mutation accumulation, DNA damage and loss of heterozygosity (LOH) in precancerous epithelium remain obscure^2,11^. This limitation creates a gap in our ability to study early transformation events *in vivo* and to fully decipher chronological and causal tumorigenic evolution in this high-risk population^2,11^. It is therefore unclear whether epithelial plasticity acts as a driver or a consequence of BRCA1-defciency associated mammary tumorigenesis^4,6,10,14^.

Here, we show that *BRCA1* heterozygosity is associated with an EMT-prone epithelial state under *ex vivo* perturbations, revealing a targetable vulnerability in patient derived organoids. EMT is a transient, multifaceted heterogeneous embryonic process that can be reactivated in response to inflammation and microenvironmental changes triggering wound-healing and tissue remodeling but also pathologic tissue fibrosis and cancer progression^15–19^. Increasing evidence in murine models was able to link EMT and inflammation induced cell-type transitions to genomic instability, mutation burden and tumor initiation in gastrointestinal tumors^12,13,20–24^. However, patient-derived early tumorigenesis models that enable the interrogation of dynamic changes remain scarce and technically challenging^2,11,16^. While somatic mutations can accumulate in normal tissue, an invasive phenotype is required for malignant progression^22,25,26^. We hypothesized that inflammation-induced mutation accumulation contemporaneous with EMT-susceptibility can drive invasive phenotypes and promote precancerous traits.

Since *BRCA1*-PV female carriers (*BRCA1* carriers) are prone to develop triple-negative breast cancer with a basal-like phenotype^27^, dissecting the role of EMT and epithelial plasticity in early tumorigenesis can provide opportunities for targeted cancer prevention in this population of high-risk patients^16^. To phenotypically delineate precancer traits in mammary epithelium of heterozygous *BRCA1* carriers, we generated patient-derived normal (non-transformed) mammary organoids (PDO) from *BRCA1* carriers and non-carriers^28^. Using external inflammatory and stress inducing stimuli (e.g. cytokines and chemotherapy), we investigate EMT, lineage plasticity, and emergent vulnerabilities in normal mammary epithelium. We identify candidate EMT-primed and EMT-induced programs enriched in subsets of *BRCA1* carriers.

### Heterozygous *BRCA1*-PV predispose mammary epithelium to EMT

We generated PDOs from 5 *BRCA1* carriers and 6 non-carriers (*BRCA1/2*-wild type (WT)) from a heterogeneous patient population spanning a range of ages (32 – 85), parity and family history of cancer (Fig. 1a, Table 1 and Supplementary Table 1). While organoid culture conditions are designed to maintain epithelial hemostasis and heterogeneity^28–31^, the altered *ex vivo* microenvironment often challenges the long-term maintenance of PDOs, particularly when derived from normal tissue. Notably, *BRCA1*-deficient normal PDOs exhibited a significantly higher rate of successful long-term establishment and expansion compared to non-carrier PDOs (Fig. 1a). PDO cultures from mammary epithelium remain challenging; the long-term maintenance rates are extremely low for WT organoids, necessitating early experimentation in primary cultures. Additionally, slow-growing cultures often limit large-scale experiments and cell-cell adhesions hinder organoid dissociation, while dissociation-induced stress can compromise analysis of stress-related processes such as EMT. However, this unique non-transformed epithelial model from a diverse patient cohort (Table 1 and Supplementary Table 1) enables investigation of dynamic transitions in normal mammary tissue while preserving heterogeneous responses. We therefore proceeded to further developed this model with these limitations in mind.

**Figure 1:**
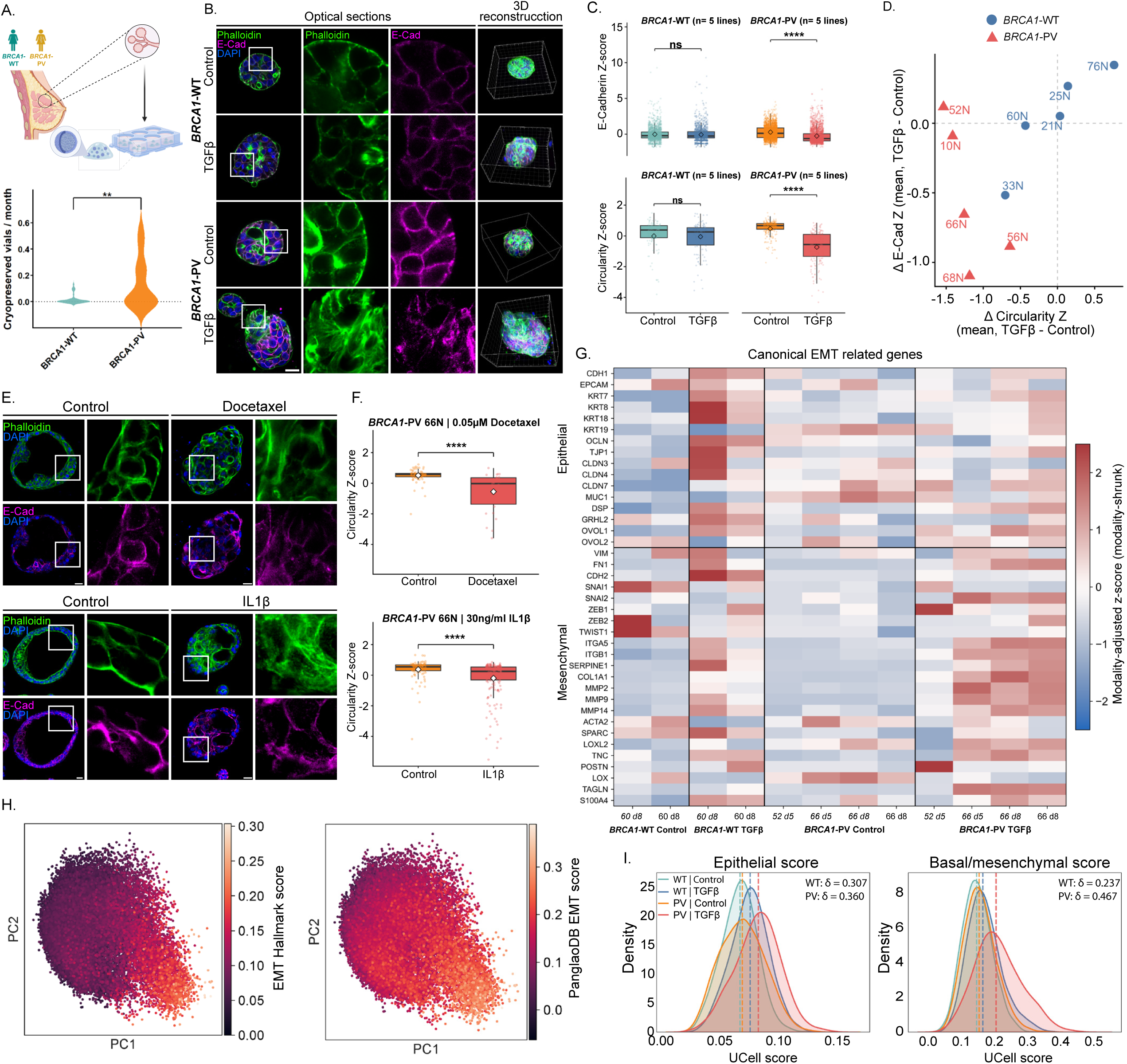
*BRCA1*-PV normal mammary gland organoids display increased EMT-susceptibility. A. Top: Establishment of patient-derived mammary organoids (PDOs) from *BRCA*-WT and *BRCA1*-PV carriers. Organoids were embedded in BME and cultured under optimized conditions. Bottom: Violin plots showing cryopreservation rate (vials per months from establishment to data collection) in *BRCA1*-WT (n = 30) and *BRCA1*-PV (n = 12) PDO lines. P value derived from a quasi-Poisson generalized linear model (*P* = 0.0011). B. 3D confocal images of organoids from *BRCA1*-WT (BR21N) and *BRCA1*-PV (BR10N) stained for E-cadherin (magenta) and F-actin (phalloidin, green) under control and TGFβ conditions. Enlargements of the squared areas are shown for E-cadherin and phalloidin stains. DAPI stain is in blue. Bar=20µm. C. Box plots of E-cadherin intensity (top) and organoid circularity (bottom) in patient-derived organoids under control and TGFβ conditions. Values were Z-score normalized within each Line × biological replicate block to account for experimental variability. Box plots show the median (center line), interquartile range (box limits), and whiskers extending to 1.5 × the interquartile range. Data points beyond this range were considered outliers and included in all statistical analyses. Each point represents a single cell (E-cadherin) or a single organoid (circularity). Error bars indicate mean ± SEM across PDO lines. D. Scatter plot of ΔCircularity versus ΔE-cadherin intensity per line following TGFβ treatment. E. *BRCA1*-PV (BR66N) PDO stained for E-cadherin (magenta) and F-actin (Phalloidin, green). Top: Control vs docetaxel (0.05µM). Bottom: Control vs IL1β (30ng/ml). Enlargements of the squared areas are shown for E-cadherin and phalloidin stains. DAPI stain is in blue. Bars=20µm. F. Single-organoid circularity in line BR66N following docetaxel (top) or IL1β (bottom) treatment. Values were Z-score normalized within biological replicate. Points represent individual organoids; diamonds indicate the mean. Statistical significance was assessed using linear mixed-effects models. n=3 biological replicates. G. Heatmap of combined single-cell and single-nucleus transcriptomic profiles showing expression levels of canonical epithelial and mesenchymal genes across genotypes and conditions. Values represent z-score normalized expression values. H. PCA plots representation of the single-cell and single-nucleus data. Left: EMT Hallmark score. Right: PanglaoDB-derived EMT score. I. Density plots of epithelial and basal/mesenchymal UCell scores (see Methods), where higher scores indicate relative upregulation of the corresponding gene signatures; dashed lines indicate medians. Cliff’s δ is shown for control vs TGFβ within *BRCA1* groups; all P < 0.001 (two-sided Mann–Whitney, BH-corrected).

**Table 1.**
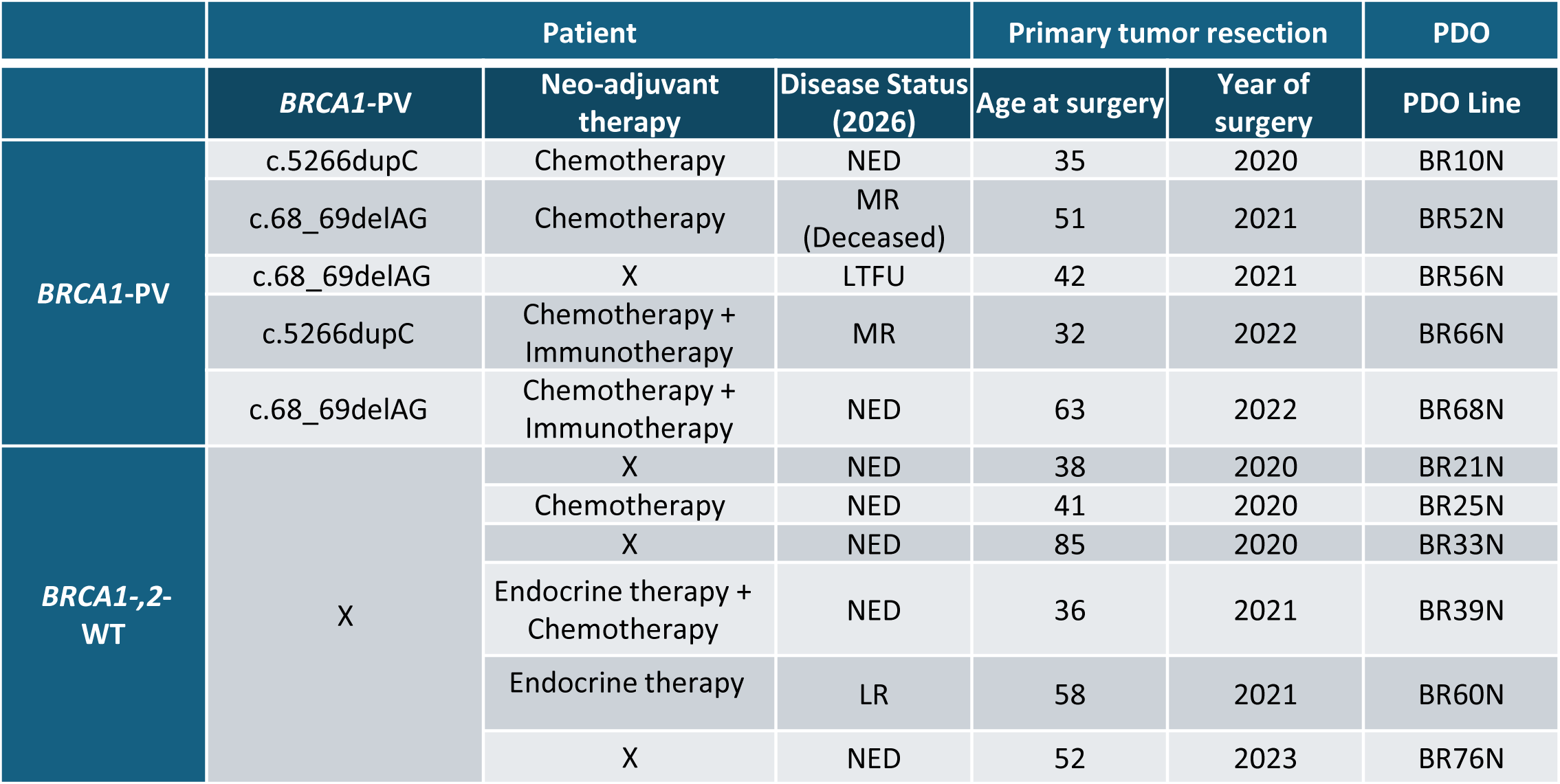
Clinicopathological and organoid characteristics of patient derived organoids. PV = Pathogenic Variant , WT = Wild Type, NED = No evidence of disease, MR = Metastatic recurrence, LTFU = Lost to follow-up, LR = Local recurrence, PDO = patient-derived organoids.

|  | Patient |  |  | Primary tumor resection |  | PDO |
| --- | --- | --- | --- | --- | --- | --- |
|  | *BRCA1*-PV | Neo-adjuvant therapy | Disease Status (2026) | Age at surgery | Year of surgery | PDO Line |
| *BRCA1*-PV | c.5266dupC | Chemotherapy | NED | 35 | 2020 | BR10N |
|  | c.68_69delAG | Chemotherapy | MR (Deceased) | 51 | 2021 | BR52N |
|  | c.68_69delAG | X | LTFU | 42 | 2021 | BR56N |
|  | c.5266dupC | Chemotherapy + Immunotherapy | MR | 32 | 2022 | BR66N |
|  | c.68_69delAG | Chemotherapy + Immunotherapy | NED | 63 | 2022 | BR68N |
| *BRCA1*-,2-WT | X | X | NED | 38 | 2020 | BR21N |
|  |  | Chemotherapy | NED | 41 | 2020 | BR25N |
|  |  | X | NED | 85 | 2020 | BR33N |
|  |  | Endocrine therapy + Chemotherapy | NED | 36 | 2021 | BR39N |
|  |  | Endocrine therapy | LR | 58 | 2021 | BR60N |
|  |  | X | NED | 52 | 2023 | BR76N |

To study the susceptibility of normal mammary epithelium to phenotypic plasticity and EMT, we treated PDOs with TGFβ, a potent inducer of EMT *in vitro* and *in vivo*^16,32^. While *BRCA1/2*-WT PDOs (hereafter WT-PDOs) maintain epithelial characteristics, even following exposure to higher concentrations of TGFβ, PDOs from *BRCA1* carriers (hereafter PV-PDOs) undergo pronounced organoid morphological changes, along with heterogeneous E-cadherin downregulation and cytoskeletal rearrangements (Fig. 1b, SFig. 1–3). To evaluate heterogeneous responses to TGFβ induction, we developed an imaging-based analytic framework enabling rapid quantification of E-cadherin downregulation at single-cell resolution within intact organoids^33^. This microscopy–deep learning pipeline circumvents the need for dissociation, avoiding stress-induced artifacts and preserving the native epithelial architecture, thereby providing a robust platform for measuring heterogeneous phenotypic changes in PDOs. The image-based analysis revealed a heterogeneous and statistically significant E-cadherin downregulation only in PV-PDOs. (Fig. 1b-c, SFig. 1-2).

**Figure 2.**
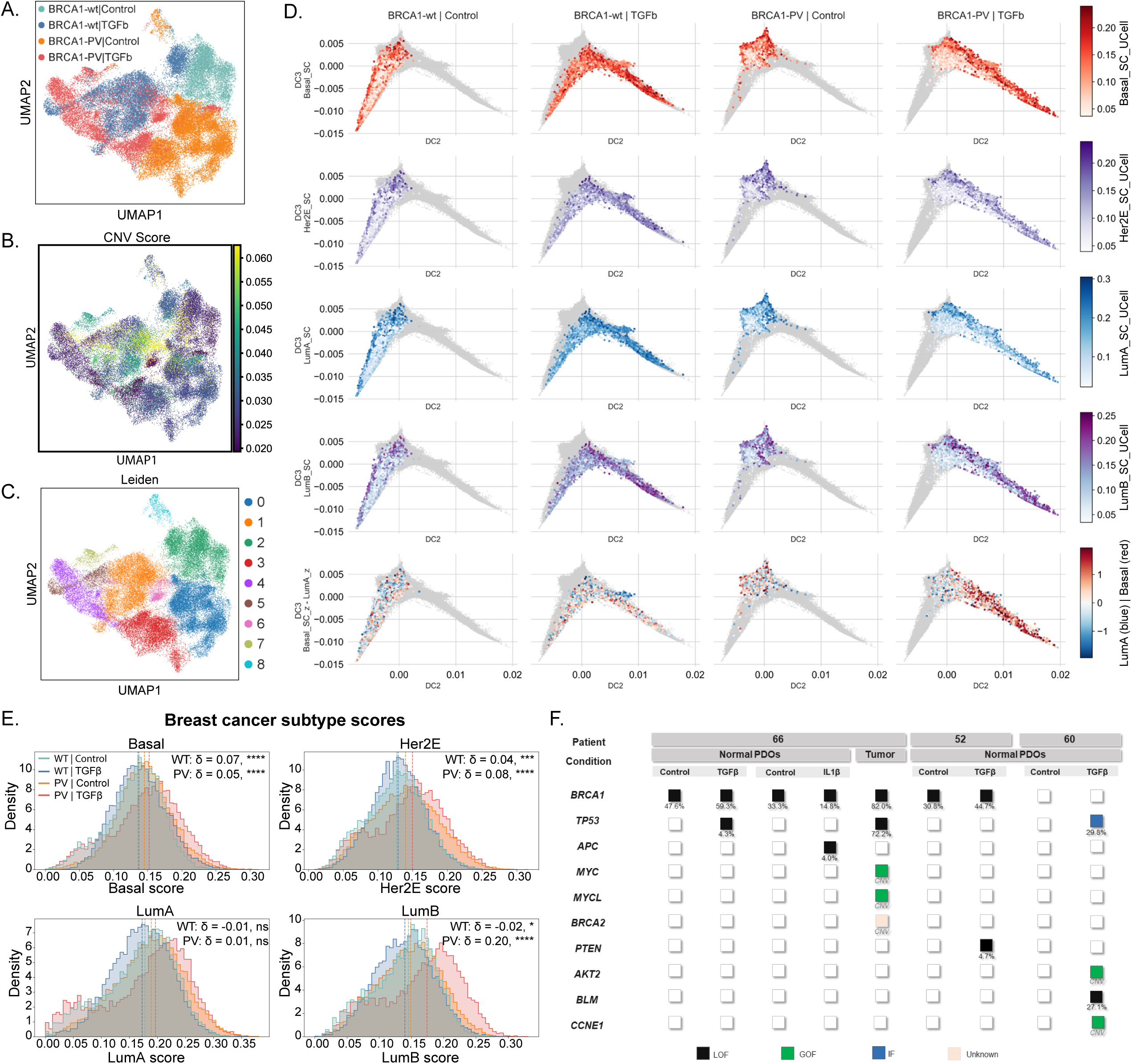
Cytokines induce enrichment of aggressive breast cancer subtypes, CNV-burden and oncogenic mutation accumulation in normal mammary organoids. A-C. UMAP projections of integrated single-cell and single-nucleus transcriptomes from patient-derived organoids (PDOs) derived from *BRCA1*-PV carriers and non-carriers under control and TGFβ conditions. A. Colored by sample annotation (genotype and treatment). B. Colored by inferred copy number variation (CNV) score. C. Colored by cluster identity, showing transcriptionally defined clusters. D. Diffusion maps plots (DC2–DC3) of Basal, HER2-positive, Luminal A, and Luminal B UCell scores in *BRCA1*-WT and PV organoids under control and TGFβ conditions. Cells from the indicated group are colored by UCell score gradient; all others are shown in gray. Bottom: Subtype balance index (Basal_SC_z − LumA_z), with positive values denoting basal-like and negative values luminal A-like enrichment. E. Histograms showing single-cell distributions of breast cancer subtype UCell scores, including basal-like (Basal), HER2-enriched (Her2E), luminal A (LumA), and luminal B (LumB) programs, where higher scores indicate relative enrichment of the corresponding gene signatures. Dashed vertical lines indicate medians. Two-sided Mann-Whitney (BH-corrected) comparing control vs TGFβ within each *BRCA1* group; Cliff’s delta. F. Genomic alterations across normal PDOs from *BRCA1*-PV lines (BR52N, BR66N), a *BRCA1*-WT line (BR60N), and the matched tumor from patient BR66N under the indicated conditions. Columns represent samples and rows represent genes. Alterations include sequence variants, annotated by variant allele frequency (VAF, %) and copy number variations (CNVs), and are classified by predicted protein functional impact (loss of function (LOF), gain of function (GOF), impaired function (IF), or unknown).

**Figure 3.**
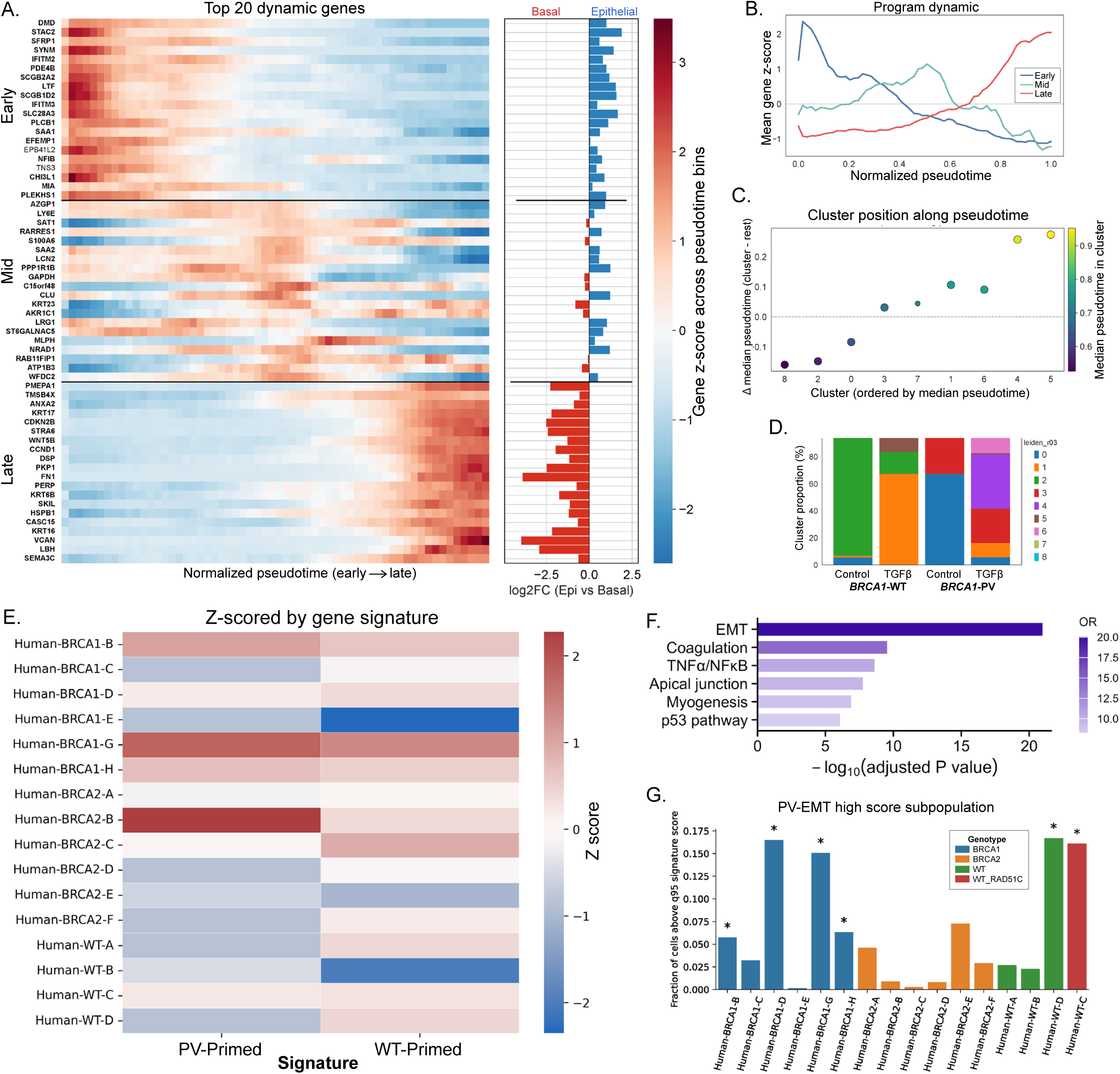
Pseudotime trajectory identifies baseline plasticity-primed and EMT-transitioning states in high-risk patients. A. Heatmap of the top 20 pseudotime-associated genes from early, mid, and late transcriptional programs, ordered along pseudotime trajectory. Cells were ordered by pseudotime, grouped into bins, and mean expression was Z-score normalized per gene. Side bars show log2 fold-change between epithelial-like and basal-like cells, defined by quartiles of the epithelial-basal axis (Epithelial_UCell − Basal_UCell). Bold gene labels indicate significant differential expression (two-sided Mann–Whitney U test, Benjamini–Hochberg FDR < 0.05). B. Line plots showing mean Z-scored expression of early, mid, and late programs along pseudotime (Normalized from 0 to 1 for visualization). C. Balloon plot of relative pseudotime position for each Leiden cluster; Color indicates median pseudotime in cluster, and dot size reflects statistical significance (two-sided Mann–Whitney test, FDR < 0.05). D. Stacked bar plot showing cluster proportions across annotation groups (*BRCA1* status and TGFβ condition). E. Heatmap of Z-scored donor-level enrichment of *BRCA1*-PV–primed and WT-primed gene signatures across donors from Gray et al. (2022)^43^. F. Top enriched Hallmark gene sets of the PV-EMT signature are shown as a horizontal bar plot ranked by −log₁₀ adjusted P value, with bar color indicating odds ratio (OR); enrichment was assessed using Fisher’s exact test with Benjamini–Hochberg correction. G. Fraction of cells per donor^43^ with high gene signature scores (≥95th percentile) for a *BRCA1*-PV–EMT signature. ∼5% are expected to exceed this threshold. Asterisks indicate significant enrichment (binomial test, FDR < 0.05).

Although EMT was induced in normal epithelial PV-PDO cultures, the resulting phenotype correlated with EMT-associated migratory capacity, demonstrating enhanced invasiveness following TGFβ induction only in organoids derived from *BRCA1*-PV carriers (Fig. 1c,d, SFig. 1-2). Next, we asked whether EMT induction was specific to TGFβ stimulation or represented a more general phenomenon of *BRCA1* carrier epithelium. PV-PDOs demonstrated a milder yet consistent EMT phenotype when treated with sub-cytotoxic concentrations of docetaxel^34^, a commonly used chemotherapy in breast cancer patients as well as the tumorigenesis-associated cytokine IL1β^35,36^ (Fig. 1e,f, SFig. 4a). Consistent with published data on extracellular stiffness induced EMT^37^, cytoskeleton rearrangements and pronounced morphological changes in organoids were also observed under stiff culture conditions (SFig. 4b).

**Figure 4.**
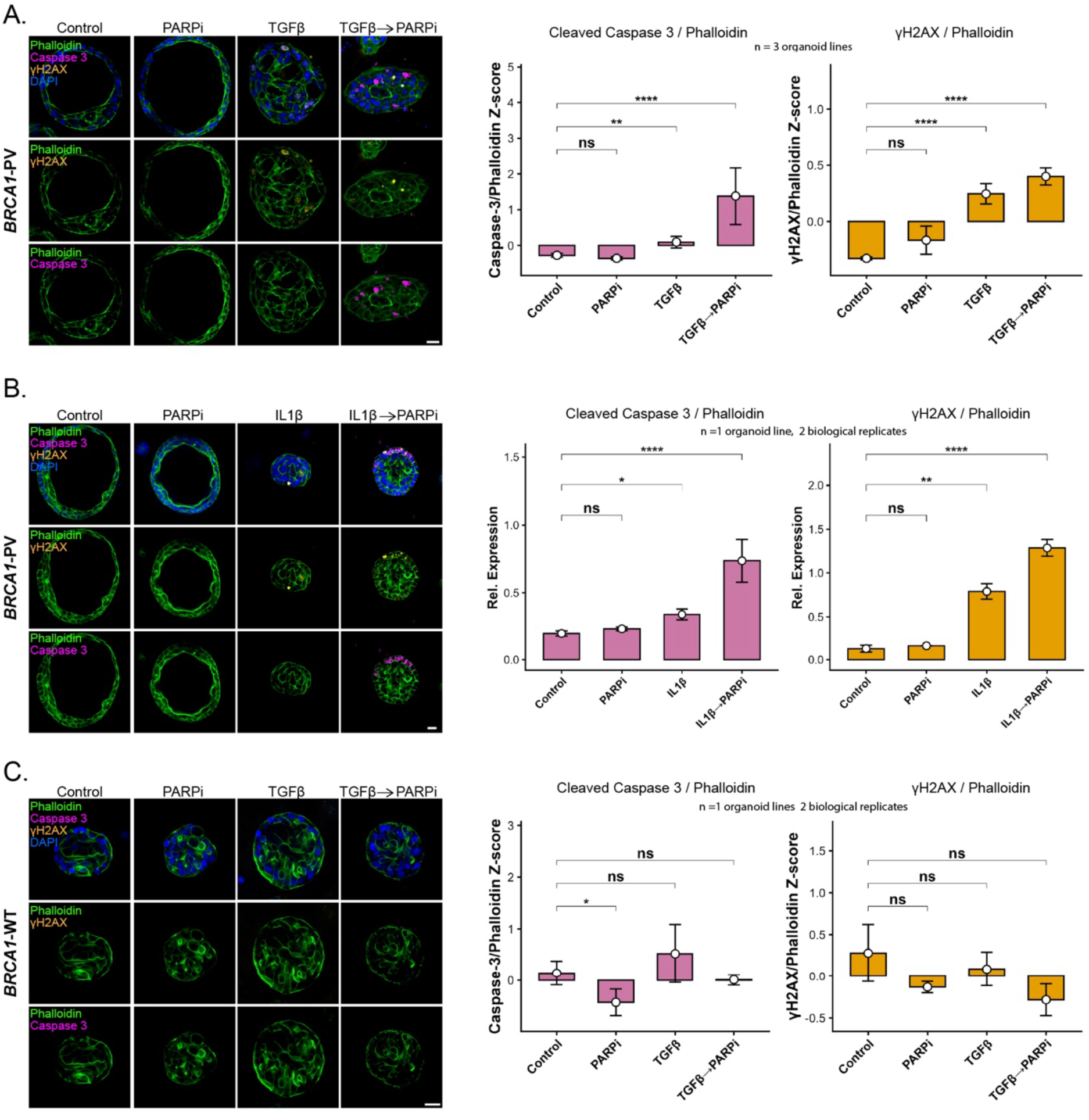
EMT induces DNA damage accumulation and PARP inhibitor susceptibility in *BRCA1*-PV mammary organoids. (A–C) PARPi response in cytokine-induced PDOs. (A) Combined analysis of three *BRCA1*-PV PDO lines (BR52N, BR66N, BR68N) following TGFβ induction. (B) *BRCA1*-PV PDOs (BR66N) following IL1β induction. (C) *BRCA1*-WT PDOs (BR39N) following TGFβ induction. Organoids were treated with DMSO (control), PARPi, cytokine alone, or cytokine followed by PARPi. Left: Representative confocal images showing phalloidin (green), γH2AX (orange), and cleaved caspase-3 (magenta). Bars=20µm. Right: Quantification of γH2AX and cleaved caspase-3 intensities measured at the single-organoid level. Marker intensities were normalized to phalloidin to account for organoid size, and Z-scored normalized for each biological replicate. For genotype-level analyses (A), values were averaged within each PDO line, and bars represent the mean across lines, with error bars indicating mean ± SEM. For single-line analysis (B,C), values were averaged within each biological replicate, and bars represent the mean across biological replicates, with error bars indicating mean ± SEM. Statistical significance was assessed using linear mixed-effects models with treatment as a fixed effect and line and biological replicate nested within line as random effects (A), or biological replicate as a random effect (B-C). Estimated marginal mean contrasts were computed versus control. ns, not significant; \**P* < 0.05; \*\**P* < 0.01; \*\*\**P* < 0.001; \*\*\*\**P* < 0.0001.

To dissect heterogeneous responses to EMT induction we performed single-cell and single-nucleus RNA sequencing (hereafter scRNA-seq). Canonical EMT markers^38^ demonstrated a robust shift toward a mesenchymal phenotype in PV-PDOs compared to WT-PDOs (Fig. 1g). Complementary EMT signatures^39,40^ were used to validate the EMT phenotype, demonstrating an enhanced global EMT-driven transcriptomic effect and a mesenchymal/basal shift in PV-PDOs treated with TGFβ (Fig. 1h,i). Together, these analyses demonstrate consistently enhanced EMT-susceptibility of PV-PDOs compared to WT-PDOs (Fig. 1, SFig. 1-4, 5a).

### EMT induces CNV burden, oncogenic mutations accumulation and a basal phenotype

Inflammatory responses have previously been linked to oncogenic mutations accumulation^23^. We therefore asked whether cytokine-induced EMT is linked to genomic instability. Indeed, scRNA-seq data predicted an increased CNV-burden in TGFβ treated organoids (Fig. 2a-c). Additionally, PV-PDOs demonstrated enhanced enrichment of breast cancer-related signatures (Fig. 2d,e)^41^.

We were intrigued to test whether oncogenic mutations could be detected in cytokines-induced PDOs and therefore performed bulk next-generation sequencing (NGS), despite anticipating that such variants would fall below <5%. We preformed TruSight Oncology 500 HR-deficiency (TSO500-HRD) profiling to detect mutations and genomic instability scores. Of note, all control PDO lines tested were negative for mutations, excluding known germline *BRCA1* variants, providing additional evidence that organoids were derived from normal mammary epithelium (Fig. 2f). However, sequencing depth was limited for all organoids and consequently changes in the relative proportion of *BRCA1* mutations should be interpreted with cautious. Despite the low sequencing depth, the results demonstrated acquired mutations in all tested PDOs following 5-8 days of TGFβ or IL1β induction. Oncogenic mutations were detected in both WT-PDOs and PV-PDOs. Notably, PV-PDO 66N line acquired TP53 loss-of-function (LOF) mutation resulting in truncated P53 protein, same as the LOF TP53 mutation acquired in the tumor specimen from the same patient (Fig. 2f). This finding is consistent with published data showing that *BRCA1* carriers commonly develop tumors harboring LOF TP53 mutations, unlike non-carriers more frequently acquiring missense TP53 mutations^42^. The TP53 mutations detected in TGFβ-induced PDO 66N and in the tumor specimen, p.R196 and p.P190Lfs57, respectively, occur in close genomic proximity, suggesting primed TP53 regions in BRCA1 carriers, as previously described in murine epigenetic studies^12^. No increase in genomic instability score was detected following induction with TGFβ or IL1β across organoid samples (Supplementary Table 2).

While cytokines exposure resulted in increased CNV-burden and oncogenic mutation accumulation, the EMT phenotype and the enrichment of breast cancer signatures were consistently more pronounced in PV-PDOs (Fig. 1-2). We therefore investigated the distribution of previously defined mammary epithelial subpopulations in WT and PV-PDOs before and after TGFβ induction^43^. While alveolar signature enrichment remained unchanged across conditions, the hormone sensing (HS) signature was enriched in both WT and PV-PDOs following TGFβ treatment. PV-PDOs treated with TGFβ showed a slight shift towards a basal signature supported by changes in cytokeratin expression (SFig. 5b-f). These subtle differences became clearer upon mapping the extended subtype signatures defined by Gray et al^43^. The HS⍺ subtype was specifically enriched in WT-PDOs, while the alveolar progenitors (AP) signature was enriched in both WT and PV-PDOs. Interestingly, higher scores for basal-luminal alveolar (BL) as well as the two myoepithelial (BA⍺ and BAβ) subtype signatures were observed following TGFβ induction, while all three showing a significant increase in PV-PDOs (SFig. 6a). These findings indicate a precancerous evolution following EMT induction in heterozygous *BRCA1* mammary organoids.

### Epithelial pseudotime progression reveals distinct *BRCA1*-PV gene signatures

To date, no *BRCA1* carrier-specific high-risk subpopulations have been described in human normal mammary epithelium^43^, although prior studies have suggested an enrichment of aberrant luminal progenitors giving rise to basal-like tumorigenesis^6,10,14^. Principal component analysis (PCA) and unbiased clustering revealed a mesenchymal/basal trajectory and the emergence of distinct transcriptional clusters in both WT- and PV-PDOs following TGFβ treatment (Fig. 1h, Fig. 2c, SFig. 7a-d). Pseudotime analysis demonstrated an early - epithelial, mid - intermediate and late - mesenchymal progression (Fig. 3a-b). Cluster distribution along the pseudotime trajectory, across conditions and genotypes, highlighted PV-PDOs specific subpopulations (Fig. 3c-d, SFig. 7a). Consistent with published data^43^, BL subtype exhibited an intermediate phenotype within the EMT trajectory (SFig. 6b). In untreated WT-PDOs we observed a nearly-monotonic dominance of an epithelial cluster #2 while untreated PV-PDOs were dominated by two intermediate clusters #0 and #3. While clusters #2 and #0 declined following TGFβ treatment in WT- and PV-PDOs respectively, cluster #3 in PV-PDOs remained largely unchanged (Fig. 3d). Clusters #2 and #0 were therefore referred to as WT-primed and PV-primed respectively as these populations changed following TGFβ induction.

Due to the challenges in expanding normal mammary PDOs described above, scRNA-seq data were generated from only one WT-PDO line (derived from a high-risk patient; see Supplementary Table 1) and two PV-PDO lines (Fig. 1g, SFig. 5a). Additionally, organoid cultures require a degree of epithelial plasticity when adapting to primary culture conditions (Fig. 1a). These model-specific limitations may lead to artificial or donor-specific transcriptomic profiles. We therefore asked whether the two primed cluster-signatures will be enriched in published scRNA-seq datasets from *BRCA* carriers and non-carriers^43^. While the WT-primed signature showed a non-specific enrichment pattern across all donors, the PV-primed signature demonstrated *BRCA*-carriers specific enrichment (Fig. 3e). However, two *BRCA1*-PV donors (*BRCA1*-C and *BRCA1*-E) showed low PV-primed signature enrichment, whereas one WT donor (WT-C) exhibited higher enrichment. Metadata from Gray et al.^43^ clarified these findings; *BRCA1*-C and *BRCA1*-E are healthy donors from the *BRCA1* cohort (no history of breast cancer), whereas WT-C is a healthy donor carrying a germline *RAD51* PV (a member of HR pathway^2^). To conclude, we utilized the transcriptomic EMT trajectory in organoids to identify PV-specific gene signature enriched in high-risk patients.

While control PDOs exhibited a relatively homogeneous cell population, TGFβ induced PDOs displayed a more heterogeneous cluster distribution, most prominently in PV-PDO (Fig. 3d). Clusters #4, #5 and #6 emerged following TGFβ treatment, while clusters #4 and #6 were unique to PV-PDOs (Fig. 3d, SFig. 7a). By comparing differences between these clusters (see Methods) we generated a TGFβ treated PV specific signature, hereafter referred to as the PV-EMT signature. PV-EMT signature exhibited statistically significant enrichment for EMT hallmark, supporting the PV-PDOs observed phenotype (Fig. 3f and Supplementary Table 3-5). We then asked whether specific donors harbor subpopulations enriched for the PV-EMT signature, potentially reflecting EMT dynamics in normal tissue. Indeed, donors showing enrichment of the PV-primed signature also exhibited significant enrichment of the PV-EMT signature, with the addition of one WT donor (Fig. 3g). To further validate this observation, we performed the same analysis in *BRCA1* and WT donors from Reed et al^44^. While neither WT donor showed enrichment, 5 of 12 *BRCA1* carriers exhibited enrichment of the PV-EMT signature (SFig. 7e).

These findings indicate that epithelial states with enhanced plasticity can be detected in specific *BRCA1* carriers, potentially with higher risk to develop breast cancer (Fig. 3e,g). EMT dynamics can be detected in the mammary epithelium of *BRCA1* carriers across independent published cohorts (Fig. 3g, SFig. 7e), supporting *in vivo* relevance. Together, these analyses define a baseline EMT-primed state and an induced EMT program that marks a high-risk epithelial trajectory in *BRCA1* carriers.

### Precancerous EMT-induced DNA-damage in *BRCA1*-PV carriers is targetable with PARPi

Poly(adenosine diphosphate-ribose) polymerase inhibitors (PARPi) cause synthetic lethality in cells harboring dysfunctional BRCA1/2, namely cells from *BRCA1* carriers that have undergone LOH resulting in impaired HR^11,45,46^. Accordingly, PV-PDOs are not expected to be affected by PARPi. In our PV-PDOs model, a small percentage of cells undergo precancerous changes, acquiring oncogenic mutations (<5% according to NGS profiling; see Fig. 2f) and adopting an invasive, basal phenotype (Fig. 1–3). We therefore hypothesized that these rare precancerous events are accompanied by DNA damage and consequently confer PARPi sensitivity. To test this hypothesis, PV-PDOs were induced to undergo EMT using either TGFβ or IL1β followed by short treatment with PARPi (Olaparib). As expected, PARPi-only treated PV-PDOs showed no increase in DNA damage or apoptosis, and WT-PDOs showed neither DNA damage following TGFβ nor response to PARPi. DNA damage accumulation was significantly increased in TGFβ or IL1β EMT induced organoids, accompanied by a significant increase in apoptosis following PARP inhibition (Fig. 4, SFig. 8). The results indicate that in response to cytokines exposure, normal mammary epithelial cells in PV-PDOs can acquire genomic instability resulting in sensitivity to short-term PARP inhibition.

## Discussion

In this study we established a non-transformed human mammary epithelium organoid model to study EMT-susceptibility in *BRCA1* carriers. Patient-derived normal *BRCA1* heterozygous mammary organoid cultures allow turning the lens on the effects of EMT in driving precancerous states. Our results indicate that *BRCA1* heterozygosity drives inherent mammary epithelial plasticity, exposing normal epithelium to increased DNA damage, accumulation of BRCA-associated oncogenic mutations, and acquisition of an invasive phenotype in response to cytokines exposure. The dynamic changes observed in organoids reflect high-risk states in mammary epithelium from *BRCA1* carriers demonstrating the relevance of the PDO model in understanding human disease trajectories.

While murine models provide evidence to the role of EMT in tumor initiation^6,10,14,21,24,47^, the possibility to study precancer events in *BRCA1* heterozygous mice is limited without the conditional activation of an additional mutation^11,12^. Recent and ongoing efforts shed light on the high prevalence of somatic mutations in non-cancerous human tissues^22,25,26^. In parallel, studies conducted with murine models were able to demonstrate the effects of inflammation in inducing genomic alterations^21,23,24^. However, identifying EMT phenotypes and deciphering the dynamic changes induced by cytokine exposure in normal human epithelium remains challenging. Particularly when these changes occur in a small cell population within the tissue^22,26^.

Modeling EMT in PDO cultures enabled the chronological dissection of tumor-initiating events in the mammary epithelium of *BRCA1* carriers. We show that, in this context, mutation accumulation following inflammatory signals is closely linked to epithelial plasticity and transition to an invasive state. Furthermore, both primed and EMT-induced gene signatures were enriched in high-risk patient datasets, supporting the potential for validation and patient stratification in larger cohorts. Finally, identifying EMT as a trigger of precancerous events in *BRCA1* carriers highlights a potential window for preventive intervention using short-term PARPi treatment.

The proposed model facilitates epithelium perturbation resulting in EMT-induced precancer traits but does not imply full tumorigenesis that would be accompanied by increased proliferation and clonal expansion. Other technical limitations of mammary PDO cultures are discussed throughout the manuscript. It is important to emphasize that while primary organoids can be established from surgical specimen in most cases, expanding and maintaining normal mammary PDOs in culture creates an experimental bottleneck. Interestingly, the most expandable WT-PDO line originated from a patient with a history of breast cancer and adjuvant endocrine therapy, suggesting a potential culture advantage for high-risk patients. Nevertheless, we demonstrate the feasibility of studying rare precancerous events and dynamic cellular plasticity in WT- and PV-PDOs. Taking the model’s limitations into account, we maintained a cohort size comparable to previous studies of normal mammary gland tissues from donors^43^ for all phenotypic studies. However, due to limited expansion potential expected in non-transformed epithelium, genomic and single-cell transcriptomic analyses were performed on a smaller cohort (1 WT-PDO line and 2 PV-PDO lines). Utilizing published datasets with extensive metadata annotation^43,44^ enabled validation of our *ex vivo* culture models for both WT- and PV-PDOs in external cohorts.

We demonstrate that normal mammary gland organoids from *BRCA1* carriers and non-carriers can be used to delineate the chronological sequence of precancerous events in this high-risk population. Importantly, our bulk TSO-500HRD analysis revealed oncogenic mutation accumulation but was not sensitive enough to detect genomic instability variations, which, based on our image-based analysis, are expected in only ∼5% of the cells. While genomic instability and LOH are necessary steps in BRCA1-related tumor initiation, dynamic perturbation of PDOs underscores EMT as a key driver of precancerous events. This in turn, reveals epithelial vulnerabilities and potential prevention strategies. Together, our data suggest that identifying EMT-associated states in high-risk patients can be used for patient stratification and may provide a strategy for cancer prevention in *BRCA1* carriers.

## Acknowledgments

We gratefully acknowledge the teams who facilitated tissue collection for these studies, including G. Hout-Siloni from the Sheba Biobank and the Pathology institute team led by I. Barshak. We thank O. Hochleitner from the Genomic facility at the MDC, Berlin for snRNA processing and profiling. We are grateful to Y. Shav-Tal and his team from the Bar-Ilan University for the support in wide-field microscopy imaging. We thank I. Shoval from Bar-Ilan University for assistance with image processing. We are thankful for the fruitful discussions with K. Lisek from the MDC in Berlin as well as E. N. Gal-Yam and M. Dadiani from the Sheba Medical Center, Israel. This work was supported by THE ISRAEL SCIENCE FOUNDATION (grant no. 2777/22), the Israel Cancer Association (grant No. 20240977) and the Basser Center for BRCA External Research Grant Program 2024. Additional support was provided by the Yoran Institute PhD Scholarship for advanced PhD students and postdoctoral fellows in personalized medicine and gene therapy. The funders had no role in study design; data collection, analysis, or interpretation; manuscript preparation; or the decision to submit for publication. Illustration in Fig. 1a was created in https://BioRender.com.

## Contributions

N.B.H, supported by R.B.Y, designed the experiments, analyzed the data and interpreted the results. The experiments were conducted by N.B.H, with assistance from R.B.Y and S.A.Y Onco-genetic interpretation and guidance were provided by R.B.M. T.M, N.H, V.N, A.M performed tissue collection and initial processing after surgery. O.G provided oncological guidance and patient selection. N.B.L and G.G performed pathological reviews. The scRNA-seq profiling were carried out by I.E.M and R.B.Y supported by the laboratory of Y.E.A. The snRNA-seq profiling were carried out by the group of T.C. Computational analysis was done by N.B.H, E.M and D.I.R supported and guided by Y.E.A and N.R. NGS and GIS profiling and analysis were conducted by P.H and Y.Z. The manuscript was prepared by N.B.H, R.B.Y and D.I.R with input from all authors. R.B. and D.I.R. provided funding for the project. D.I.R. conceived the project, provided guidance in experimental design and analysis.

## Materials and Methods

### Mammary Gland Patient Derived Organoids Cultures

For the establishment of patient-derived mammary gland organoids, normal breast tissues were obtained via the Sheba Tissue Bank from patients that underwent mastectomy or lumpectomy under informed consent. Following the published protocol^31^ the tissue was both mechanically and enzymatically digested, and isolated breast cells were plated in adherent Cultrex growth-factor-reduced basement membrane extract (BME) type 2 drops ((R&D Systems; 3533-005-02)). Organoids were overlaid with optimized organoid culture medium^31^ containing 10% R-spondin-1-conditioned medium (RCM) produced from HEK293 HA–Rspo1–Fc cells (Cultrex® HA–Rspondin-1–Fc 293T cells; 3710-001-01), 20% Wnt3a-conditioned medium (WCM) produced from L cells stably transfected with pcDNA3.1-Zeo-mouse Wnt3a, and 10% Noggin-conditioned medium (NCM) produced from HEK293 cells stably transfected with pcDNA3-mouse NEO insert (to confer neomycin resistance; cells for WCM and NCM production were kindly provided by the Hubrecht Institute, Utrecht, The Netherlands), 500nM A83-01 (Tocris Bioscience; 2939), and 5 mM Y-27632 (ROCK inhibitor, Sigma Aldrich; Y0503) which is only required when organoids are first established or expended. Medium was changed every 4 days, and organoids were passaged every few weeks using mechanical shearing using TrypLE Express (Invitrogen; 12605036). All Organoids lines were grown at 37^0^C, 5% CO2, 95% humidity.

Use of human tissues was approved by the local ethics committee and by the Associate Director at the Sheba Medical Centre (approval no. 7188-20-smc), and informed consent was obtained for all tissue donors. Investigations were conducted according to the principles expressed in the Declaration of Helsinki.

### Clinical data collection

Clinical data were obtained from institutional medical records and collected under institutional review board approval, in accordance with the Declaration of Helsinki. All Patients follow-ups were updated as of April 2026.

### EMT induction in patient-derived organoids

Whole organoids were embedded in BME (R&D Systems; #3533-010-02) and plated in 18-well μ-Slides (ibidi; #81816). Organoids were cultured for 3–10 days, with medium refreshed every 3 days.

#### TGFβ

Control organoids were maintained in standard organoid culture medium. For EMT induction, organoids were cultured for 8-10 days in EMT medium defined as organoid culture medium lacking A83-01 (a TGFβ signaling inhibitor) and supplemented with 2 ng/ml recombinant TGFβ (R&D Systems; #240-B), as we previously described^31^. In experiments assessing the effect of A83-01 withdrawal alone, organoids were cultured in medium lacking A83-01 without TGFβ supplementation.

#### IL1β

Organoids were cultured for 8 days in either standard medium (Control) or medium supplemented with recombinant IL1β (R&D Systems; #201-LB) (30ng/ml)

#### Docetaxel

Organoids were cultured for 8 days in either standard medium (Control) or medium supplemented with Docetaxel (0.05µM).

#### Matrix stiffness

Matrix stiffness protocol was adapted from a previously described method^48^. Briefly, 8-well µ-Slides (ibidi; 80806) were coated with a 1:1 mixture of growth medium and either stiff (3 mg/ml collagen) or soft (1.2 mg/ml collagen) ECM. ECM is composed of either stiff (3 mg/mL) or soft (1.2 mg/mL) collagen. The matrix was prepared using Matrigel Basement Membrane Matrix (Corning; 354234) and Collagen Type I, Rat Tail (Corning;354236). Organoids were resuspended in a 1:1 mixture of growth medium and ECM, then seeded onto the pre-coted gels. After polymerization, culture medium was added and replaced every two days.

### PARPi treatment following EMT induction in patient derived organoids

Following a 4-day induction with TGFβ or IL1β as described above, organoids were treated for 24 h with the PARP inhibitor Olaparib (MedChemExpress; HY-10162) at a final concentration of 5 µM. Control samples were treated with dimethyl sulfoxide (DMSO; Sigma-Aldrich; D2653).

### Immunofluorescence staining

Following treatments, organoids were fixed in 4% paraformaldehyde (PFA; Bio-Lab; #0006450323F1) for 30 min, permeabilized with 0.3% Triton X-100 in PBS for 30 min and blocked in 3% bovine serum albumin (BSA; Avantor; 0332-50G) in PBST (0.01% Triton X-100 in PBS) for 1h.

For E-cadherin, cleaved Caspase-3, γH2AX, and F-actin (Phalloidin) staining, organoids were incubated with primary and corresponding fluorophore-conjugated secondary antibodies as detailed in Supplementary Table 6. Primary antibody incubation was performed for 2 h at room temperature (RT), followed by secondary antibody incubation for 1h at RT, together with Phalloidin (Abcam; ab176753; 1:1000) for F-actin staining.

For cytokeratin staining, the same protocol was followed, except that directly conjugated primary antibodies were used (Supplementary Table 6).

All antibodies were diluted in 3% BSA in PBST. Each incubation step was followed by three washes in PBST (5 min each). Nuclei were counterstained with 4′,6-diamidino-2-phenylindole (DAPI; Invitrogen; D1306; 1:1000( for 10 minutes, and organoids were maintained in ibidi mounting medium (ibidi; IBD-50001) until imaging.

### FFPE Blocks

Following treatments, the organoids were extracted from BME by 1h incubation with Cell Recovery Solution (Corning®; 354253) on ice and fixed in 4% paraformaldehyde for 30 minutes. After fixation, the organoids were washed with PBS, suspended in 1% Agarose and embedded in paraffin to create formalin-fixed paraffin-embedded (FFPE) blocks. FFPE blocks were sectioned using a microtome.

#### Immunostaining

Slides were dried at 60–65 °C overnight, stored at 4 °C, and processed within one week. Antigen retrieval was performed in 1X Omniprep buffer at 90–95 °C for 1 h, followed by gradual cooling and sequential washes in distilled water and PBST (PBS + 0.05% Tween-20). Sections were blocked with 10% BSA for 1 h at room temperature and incubated overnight at 4 °C with primary antibodies Alexa Fluor 647–conjugated anti-cytokeratin 14 (CK14; Abcam; ab206100; 1:100) or Alexa Fluor 488–conjugated anti-cytokeratin 8 (CK8; Abcam; ab192467; 1:100)) diluted in 2% BSA. The following day, slides were washed 3 times with PBST, counterstained with DAPI (1:1000, 5 min), and mounted with ibidi antifade medium. Coverslips were sealed with nail polish, dried, and stored at 4 °C until imaging.

### Confocal microscopy

Confocal imaging was performed on a confocal LSM700 ZEISS microscope, using 40x air objective, NA 1.518, and on TCS SP8 Leica Confocal microscope, equipped with a Leica DFC9000 GT camera, using HC PL FLUOTAR 40x, NA 0.8 or HC PL FLUOTAR 10x, NA 0.3 air objective (Leica Microsystems, Nussloch, Germany).

### Widefield microscope

Wide-field fluorescence images were obtained using an Olympus IX83 fully motorized inverted microscope (40× UPLSAPO 2 air objective, 0.95 NA) fitted with the Prime BSI camera (Photometrics) driven by the CellSens software

### Image processing, quantification and analysis

Image processing (i.e., Crope, pseudo-coloring, merge channels) was performed with LAS X software (3.7.2.22383) and the Fiji processing software package of ImageJ (1.51n). 3D reconstruction was performed using Imaris (Bitplane).

#### Single-cell E-cadherin intensity

Single cells within confocal images of organoids were segmented using Cellpose3, a deep learning-based instance segmentation algorithm^49^. DAPI was used as a nuclear marker and phalloidin as a cytoplasmic marker to improve segmentation accuracy. Segmentation quality was visually validated against manual annotation. For each segmented cell, a binary mask was generated, and mean gray-level intensity (8-bit, 0–255) was quantified for E-cadherin and cytokeratin channels, as we previously described^33^. Single-cell E-cadherin intensities were extracted from segmentation masks. Mean fluorescence intensity per cell was calculated for each channel, and distributions were visualized using kernel density estimation, with Control and TGFβ conditions overlaid within each group (Supplementary Tables 7-8).

#### Single organoid circularity

Organoid circularity was quantified by manually outlining each organoid in the Fiji processing software package of ImageJ (1.51n). Circularity was calculated as 4π × Area / Perimeter² as previously described^50^. Circularity measurements were analysed at the single-organoid level (Supplementary Tables 9-12).

#### Joint circularity and E-cadherin analysis

For each PDO line, treatment response was defined as the difference between TGFβ and Control in mean circularity and mean E-cadherin Z-score. These organoid-line-level deltas were combined and visualized in a two-dimensional scatter plot.

#### PARPi experiments - γH2AX and cleaved Caspase-3 intensities

γH2AX and cleaved Caspase-3 intensities were quantified at the organoid level from confocal images using Leica LAS X software (Leica Microsystems, v3.3.0). Regions of interest were manually defined based on phalloidin staining outlining organoid boundaries. Mean fluorescence intensity was measured within manually defined organoid regions. To account for differences in organoid size, marker intensities were normalized to phalloidin signal (Supplementary Tables 13-14).

### ScRNA SnRNA Seq ScRNA seq

Single-cell RNA-seq libraries were prepared at the Crown Genomics Institute of the Nancy and Stephen Grand Israel National Center for Personalized Medicine, Weizmann Institute of Science, using the 10X Genomics technology. Following 10 days of culture under control or TGFβ conditions (as detailed above),organoids were resuspended in TrypLE Express (Invitrogen; 12605036) and dissociated into a single-cell suspension using mechanical shearing. Dissociated organoids were filtered with 15µm cell strainer (pluriSelect; 43-50015-01). Cells were counted, and viability was assessed using Trypan blue. Cells were diluted in PBS + 0.04% BSA to a final concentration of 1000 cells/µl and immediately processed with the Chromium Next GEM Single Cell 3’ v3.1 kit, according to the manufacturer’s protocol. Final libraries were quantified by qPCR with the NEB-next Library Quant Kit (New England Biolabs) and with Qubit and TapeStation. Sequencing was done on a Nova-Seq6000 using SP, 100 cycles kit mode, allocating 800M reads in total (Illumina).

### SnRNA seq

Following 5-8 days of culture under control or TGFβ conditions, the organoids were extracted from BME by 1h incubation with Cell Recovery Solution (Corning®; 354253) on ice and fixed in 4% paraformaldehyde for 30 minutes. After fixation, the organoids were washed with PBS, suspended in 1% Agarose and embedded in paraffin to create formalin-fixed paraffin-embedded (FFPE) blocks. FFPE blocks were sectioned to 25µm slices using a microtome.

To obtain single nuclei suspensions form FFPE tissue sections, the samples were processed according to the Sample Preparation from FFPE Tissue Sections for GEM-X Flex Gene Expression using the Xylene-based Protocol - gentleMACS (10X Genomics, CG000784 | Rev C). Isolated nuclei were counted with Countess 3 FL (Invitrogen). Single nuclei suspensions were further processed for snRNA-seq analysis according to the GEM-X Flex Gene Expression Reagent Kits for Multiplexed Samples (10X Genomics, CG000787 | Rev B). The concentration of the library was measured using a Qubit 3 Fluorometer (Qubit). The fragment size distribution was determined using a 4200 TapeStation system (Agilent Technologies). The libraries were sequenced in a NovaSeq X Plus sequencer (Illumina) according to 10X Genomics recommendations. Raw sequencing data were processed using the Cell Ranger pipeline (10x Genomics, version 7.0.0), including read alignment to the reference genome, UMI counting, and generation of gene–cell count matrices.

### Analysis

Single-cell (sc) and single-nucleus (sn) RNA sequencing (RNA-seq) data were processed using Python (v3.14.2) and *Scanpy* (v1.11.5). The dataset comprised three PDO lines: BR52N (BRCA1-PV), BR60N (BRCA1-WT), and BR66N (BRCA1-PV), spanning 12 independent experiments. BR52N was profiled exclusively using single-nucleus RNA-seq, whereas BR60N and BR66N included both single-cell and single-nucleus RNA-seq data. Cells were distributed across Control and TGFβ-treated conditions (Supplementary Table 15). Cell-level metadata were curated, including patient-derived organoid (PDO) line, treatment condition, modality (scRNA-seq or snRNA-seq), and experimental annotations. To enable integrated analysis across modalities, datasets were restricted to the intersection of shared gene. Highly variable genes were identified using the *Seurat* v3 method (n_top_genes = 2000) where the genes are shared between modalities. The integrated dataset was used for downstream dimensionality reduction, clustering, and transcriptional analyses.

Dimensionality reduction and clustering were performed in *Scanpy*^51^. Principal component analysis (PCA; 50 components) was computed on highly variable genes, and Harmony integration was applied to correct for technology-driven batch effects using modality as the integration variable. A k-nearest neighbor graph (k = 15) was constructed from Harmony-corrected embeddings and used for Leiden clustering (resolution = 0.3) and visualization. Low-dimensional embeddings were generated using UMAP and diffusion maps; UMAP embeddings were used to visualize cluster structure and sample annotations (*BRCA1* status and treatment).

UCell scores were computed as previously described^52^ using gene signatures derived from PanglaoDB^39^ Gene signatures included epithelial, mesenchymal/basal, and epithelial differentiation programs (alveolar, hormone-sensing, basal). Additional epithelial subtype signatures were obtained from the human breast atlas single-cell transcriptomics (Gray et al., 2022^43^). Breast cancer subtype-associated programs (Basal-like, HER2-positive, Luminal A, Luminal B) derived from published single-cell breast cancer atlas data (Wu et al.,2021^41^). An EMT index was computed at the single-cell level as the difference between Basal_UCell and Epithelial_UCell scores (Basal_UCell - Epithelial_UCell), where higher values indicate a more mesenchymal/basal-like transcriptional state. In parallel, EMT scores were independently computed using the Hallmark epithelial-mesenchymal transition gene set from the Molecular Signatures Database^40^, using the same UCell framework. UCell score distributions were compared across groups defined by *BRCA1* status and treatment. Distributions were visualized using normalized histograms with medians indicated by dashed lines, with normalization performed independently for each group.

Canonical epithelial and mesenchymal gene expressions were analyzed using sample-level pseudobulk profiles (Supplementary Table 16). Expression was averaged within each sample × modality, Z-score normalized across samples within each modality, and shrunk toward group means to reduce noise in low-cell-number combinations. Values were then averaged across modalities to obtain a single relative expression value per sample (z-score), and the resulting sample-by-gene matrix was visualized as a heatmap.

Copy number variation (CNV) profiles were inferred using *infercnvpy* (v0.6.1), available at github.com/icbi-lab/infercnvpy. This tool is an extension of the principles used in inferCNV^53^ with Python (v3.12.12). Gene coordinates were assigned based on GENCODE v49, and *BRCA1*-WT cells were used as reference. CNV inference was performed using a 10-gene sliding window (step size 2). A per-cell CNV burden score was defined as the mean absolute CNV deviation across genomic windows and normalized to a 0–1 scale. CNV burden was summarized at the cluster level, including mean burden, fraction of CNV-high cells (≥0.7), and cell counts.

Pseudotime trajectories were inferred using diffusion pseudotime (DPT) implemented in *Scanpy* calculated on the first 10 diffusion components, as the mesenchymal/basal program represented the dominant source of variation in principal component analysis. To minimize genotype-driven bias in root selection, basal-like cells from clusters 4 and 5 (corresponding to *BRCA1*-PV and *BRCA1*-WT populations, respectively) were used to define the starting population. For each iteration, 10 cells per cluster were randomly sampled as root cells, and DPT was computed independently. This procedure was repeated 20 times, and pseudotime values were averaged across runs to ensure robustness. Because basal-like states (populating clusters 4 and 5) emerged predominantly following TGFβ induction, pseudotime directionality was reversed to reflect the expected biological progression from epithelial to mesenchymal states. Genes associated with pseudotime were identified based on correlation with pseudotime values and grouped into early, intermediate, and late temporal programs according to their peak expression along the trajectory. For visualization, cells were binned along pseudotime, and gene expression was averaged within bins and Z-score normalized across bins. Cluster-level pseudotime position was visualized using a dot plot of the difference in median pseudotime (cluster vs. remaining cells), where point size reflects statistical significance and color encodes median pseudotime per cluster.

For gene signature derivation, WT-primed and PV-primed programs were defined by differential expressions between epithelial clusters. Genes upregulated in cluster 0 relative to cluster 2 and vice versa were used to construct cluster-specific signatures corresponding to PV-primed and WT-primed states, respectively (Supplementary Table 3,17).

The *BRCA1*-PV–associated EMT (PV-EMT) signature was derived by integrating cluster identity, treatment condition, and genotype. Genes upregulated in clusters 4 and 6 versus cluster 5 under TGFβ were intersected with genes upregulated in BRCA1-PV cells under TGFβ relative to both PV controls and *BRCA1*-WT TGFβ cells, using stringent thresholds (log2FC > 0.75, FDR < 0.01, expression fraction ≥ 10%). Intersected genes were ranked by consistency across contrasts, prioritizing genes with uniformly high effect sizes across all comparisons, and a compact signature (N = 25–200) was selected based on optimal discrimination of *BRCA1*-PV TGFβ cells using UCell scoring (AUC, median score separation, and top-decile purity). Functional enrichment was assessed using Hallmark gene sets^40^ (Supplementary Table 4,5).

To evaluate generalizability, signatures were applied to two independent human breast reference atlases (Gray et al., 2022^43^ and Reed et al., 2024^44^). External datasets were processed using a consistent *Scanpy* workflow. Only epithelial cells from both datasets were analyzed. For primed signatures, UCell scores were summarized at the donor level and Z-score normalized across donors. For PV-EMT signature, to identify donors enriched for rare PV-EMT–high subpopulations, cells exceeding a global score threshold (defined by the upper quantile of the UCell score distribution) were classified as high-scoring.

### Biostatistics

#### Cryopreserved vials

To account for variation in culture duration, freezing counts were analyzed using a generalized linear model with a quasi-Poisson distribution, including *BRCA1* status and time from establishment to data collection as covariates (Freezings ∼ Time_months + *BRCA1* status). The quasi-Poisson model was used to account for overdispersion in count data. Statistical significance for the effect of *BRCA1* status was derived from this model (Supplementary Tables 18-19).

#### Image analysis

Statistical analyses were performed in R (v4.5.1) using *lme4*, *lmerTest*, and *emmeans*. PDO lines were treated as independent biological units, and biological replicates within each line (independent experimental repeats) were modeled as nested factors. Observational units were defined according to the assay: individual cells for E-cadherin analyses, and individual organoids for circularity and PARPi experiments. Hierarchical data structure was accounted for using linear mixed-effects models where appropriate. All tests were two-sided, and *P* values were adjusted using the Benjamini–Hochberg method unless otherwise specified.

#### Data normalization

To account for experimental variability, measurements were Z-score normalized within each Line × Biological replicate block. Only finite values were included in the analysis.

#### E-cadherin and circularity analysis

For single-cell E-cadherin intensity and organoid circularity under TGF-β conditions, linear mixed-effects models were fitted separately per genotype (n = 5 lines per group) to assess treatment effects within biologically distinct groups, with condition as a fixed effect and random intercepts for line and biological replicate nested within line. For analyses restricted to a single PDO line (for example, IL1β or Docetaxel experiments), biological replicate was included as a random effect; when only one biological replicate was available, linear models were applied. Inference in these cases is restricted to within-line variation. No data points were excluded based on outlier criteria, as extreme values may reflect biologically relevant heterogeneity (SFig.1-2).

#### γH2AX and cleaved Caspase-3 analysis

For PARPi experiments, γH2AX and cleaved Caspase-3 intensities (normalized to Phalloidin to account for organoid size) were analyzed separately per genotype using linear mixed-effects models with treatment group as a fixed effect and random intercepts for line and biological replicate nested within line.

#### Statistical reporting

Estimated marginal means (EMMs) and contrasts versus Control were computed using emmeans, and effect sizes are reported as differences in EMMs (ΔEMM). Data are presented as mean ± Standard Error of the Mean (SEM). across lines for genotype-level analyses and across biological replicates for single-line analyses. Full statistical outputs, including model coefficients, adjusted P values, and sample sizes, are provided in Supplementary Table 20-25.

### scRNA and snRNA seq analysis

#### UCell score distributions

UCell score distributions were compared using two-sided Mann–Whitney U tests, with effect sizes reported as Cliff’s delta. P-values were adjusted using the Benjamini–Hochberg (BH) method (Supplementary Table 26-29).

#### Pseudotime-associated features

Gene–pseudotime associations were assessed by correlation analysis with BH correction. Differential expression between epithelial-like and basal-like populations and cluster-level pseudotime comparisons were evaluated using Mann–Whitney U tests with BH correction. (Supplementary Table 30-31).

#### Gene signatures

For primed gene signatures, analyses were performed at the donor level to avoid pseudoreplication. UCell scores were summarized per donor using the median, and differences across genotype groups were assessed using Kruskal–Wallis tests, followed by pairwise two-sided Mann–Whitney U tests. BH correction was applied within each signature. Z-score normalization was used for visualization only. For the PV-EMT signature, enrichment of high-scoring cells was assessed using one-sided binomial tests comparing observed counts to expected frequencies defined by either the quantile threshold or the global proportion of high-scoring cells. BH correction was applied, and donors with FDR < 0.05 were considered significantly enriched. (Supplementary Table 32-33).

#### Hallmark pathway enrichment

Enrichment significance was assessed using Fisher’s exact test, with P values adjusted for multiple testing using the Benjamini-Hochberg method. Hallmark terms with adjusted P < 0.05 were considered significant.

### Next Generation Sequencing

#### Ethics approval

Human tissue samples were obtained with informed consent under protocols approved by the Institutional Review Board of Sheba Medical Center, (approval no. 8621-11), in accordance with the Declaration of Helsinki.

#### Tissue processing and nucleic acid isolation

Genomic DNA (gDNA) was extracted from FFPE sections of primary patient biopsy and organoid cultures using the QIAamp DNA FFPE Tissue Kit (Qiagen), according to the manufacturer’s instructions.

#### Library preparation and hybrid-capture sequencing

Comprehensive genomic profiling was performed using the TSO500-HRD assay (Illumina). gDNA was fragmented and subjected to hybrid-capture target enrichment using a probe set covering 523 cancer-associated genes and sub-telomeric region for genomic instability score (GIS).

#### Bioinformatics pipeline and HRD scoring

Data were processed using the DRAGEN TSO500-HRD pipeline (Illumina) for alignment (GRCh37/hg19) and detection of SNVs, indels, copy number alterations, and structural variants. HRD status was determined using GIS, defined as the sum of LOH, telomeric allelic imbalance (TAI), and large-scale state transitions (LST)^54^. Standard size thresholds were applied for each metric, and a cutoff of ≥42 was used to classify HRD-positive samples^55^. Tumor-only analysis was performed using a reference-based approach to filter germline variants^55,56^. Clinical variant annotation, prioritization, and tier-based classification were performed using the Velsera PierianDx Clinical Genomics Workspace (CGW).

### Artificial Intelligence (AI) Useage

AI-assisted tools (e.g., OpenAI Codex) were used to support aspects of computational analysis. All code, analyses, and results were reviewed and validated by the authors.

## Notes

### Competing Interest Statement

The authors have declared no competing interest.

