## Supplementary Figures for "Epithelial mesenchymal transition initiates precancer states in *BRCA1* mutation carriers"

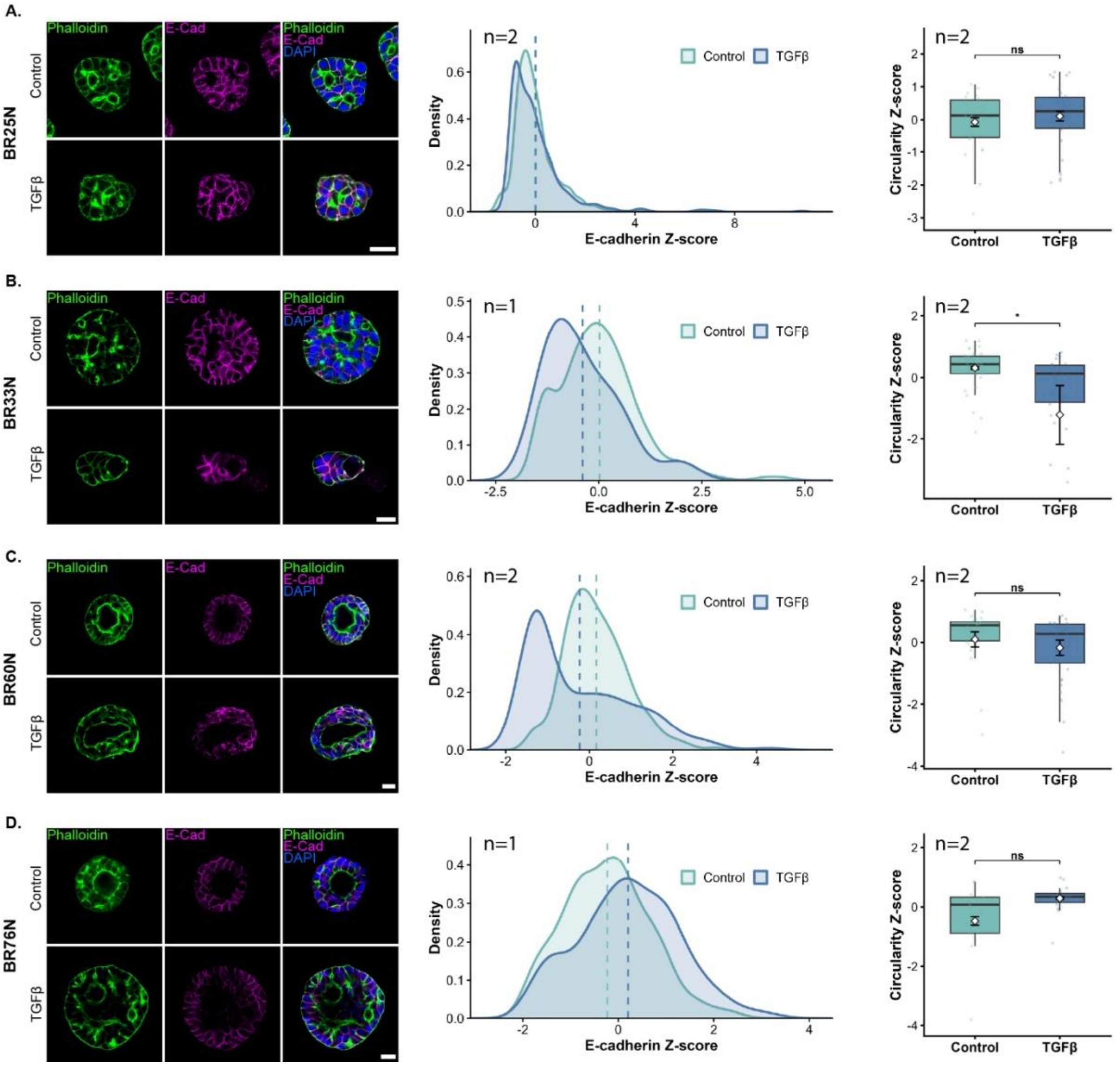


**Supplementary Figure 1. Normal mammary gland organoids from *BRCA1*-WT patients maintain E-cadherin expression and a non-invasive phenotype following TGFβ induction.**

(A–D) Confocal images and per-line quantitative analyses for individual *BRCA1*-widtype (WT) organoid lines: (A) 25N, (B) 33N, (C) 60N, and (D) 76N. Left: confocal images of organoids immunolabeled for E-cadherin (magenta), F-actin (Phalloidin, green) and DAPI (blue) under control (top) and TGFβ (bottom) conditions (Bars=20μm). Middle: density plots of single-cell E-cadherin Z-scores shown separately for each line under control and TGFβ conditions, with dashed lines indicating per-condition means. E-cadherin values were Z-score normalized within each Line × biological replicate block. Right: boxplots of single-organoid circularity Z-scores, shown per line. Circularity was Z-score normalized within each Line × biological replicate block. Each point represents one organoid; center lines indicate medians, boxes show interquartile ranges (IQR), and whiskers extend to 1.5 × IQR. Overlaid points and error bars represent mean ± SEM across biological replicates. Statistical significance was assessed per line using linear mixed-effects models (Circularity Z ~ Condition + (1|biological replicate)) with two-sided testing. ns, not significant; *P < 0.05; **P < 0.01; ***P < 0.001; ****P < 0.0001. n= biological replicates.

### Supplementary Figures

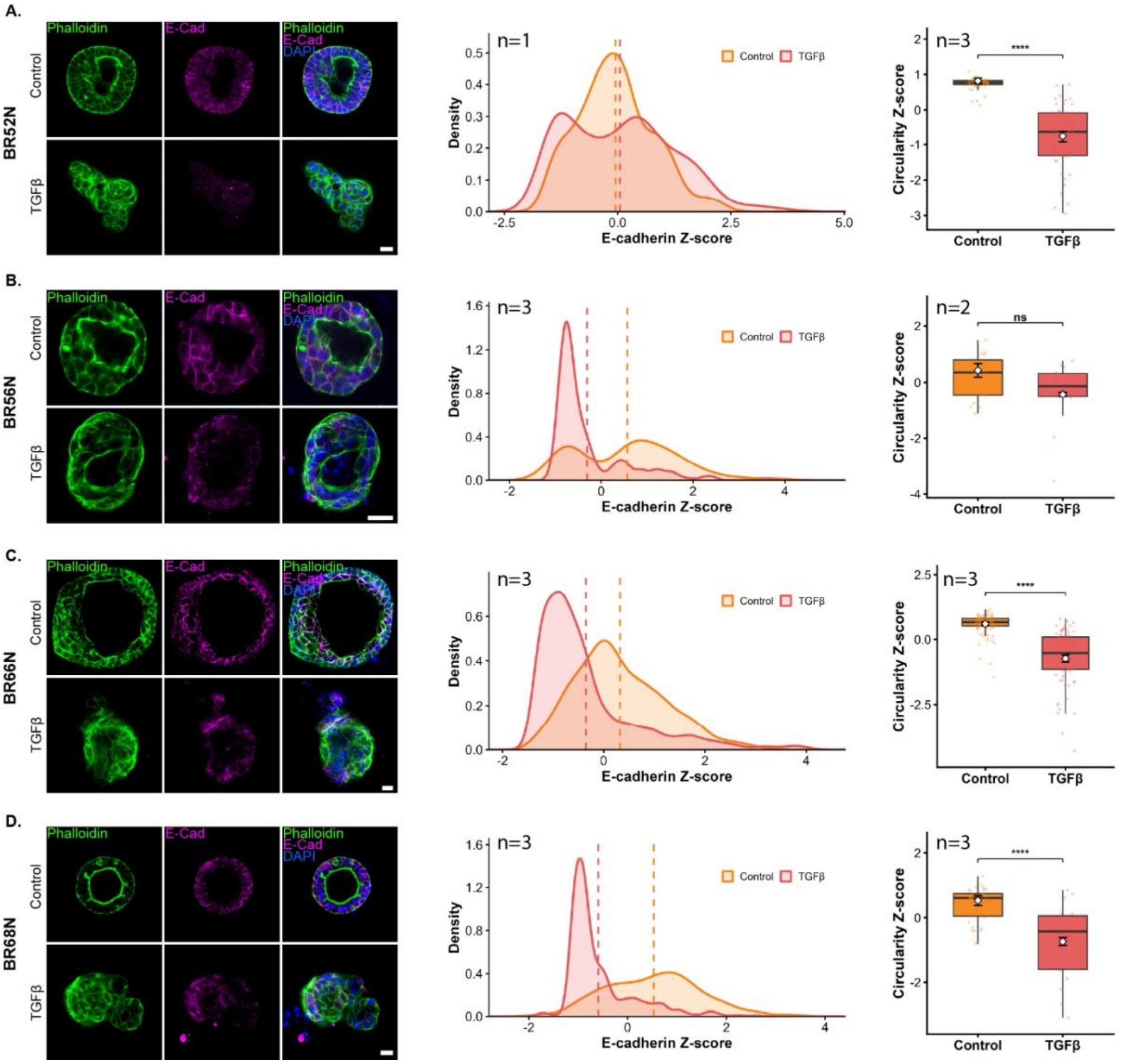


**Supplementary Figure 2. TGFβ induces heterogeneous E-cadherin downregulation and an invasive phenotype in *BRCA1*-PV normal mammary organoids.** (A–D) Confocal images and per-line quantitative analyses for individual *BRCA1*-PV organoid lines: (A) 52N, (B) 56N, (C) 66N, and (D) 68N. Left: confocal images of organoids immunolabeled for E-cadherin (magenta), F-actin (Phalloidin, green) and DAPI (blue) under control (top) and TGFβ (bottom) conditions (Bars=20μm). Middle: density plots of single-cell E-cadherin Z-scores shown separately for each line under control and TGFβ conditions, with dashed lines indicating per-condition means. E-cadherin values were Z-score normalized within each Line × biological replicate block. Right: boxplots of single-organoid circularity Z-scores, shown per line. Circularity was Z-score normalized within each Line × biological replicate block. Each point represents one organoid; center lines indicate medians, boxes show interquartile ranges (IQR), and whiskers extend to 1.5 × IQR. Overlaid points and error bars represent mean ± SEM across biological replicates. Statistical significance was assessed per line using linear mixed-effects models (Circularity_Z ~ Condition + (1|biological replicate)) with two-sided testing. ns, not significant; *$P < 0.05$; **$P < 0.01$; ***$P < 0.001$; ****$P < 0.0001$. n= biological replicates.

### Supplementary Figures

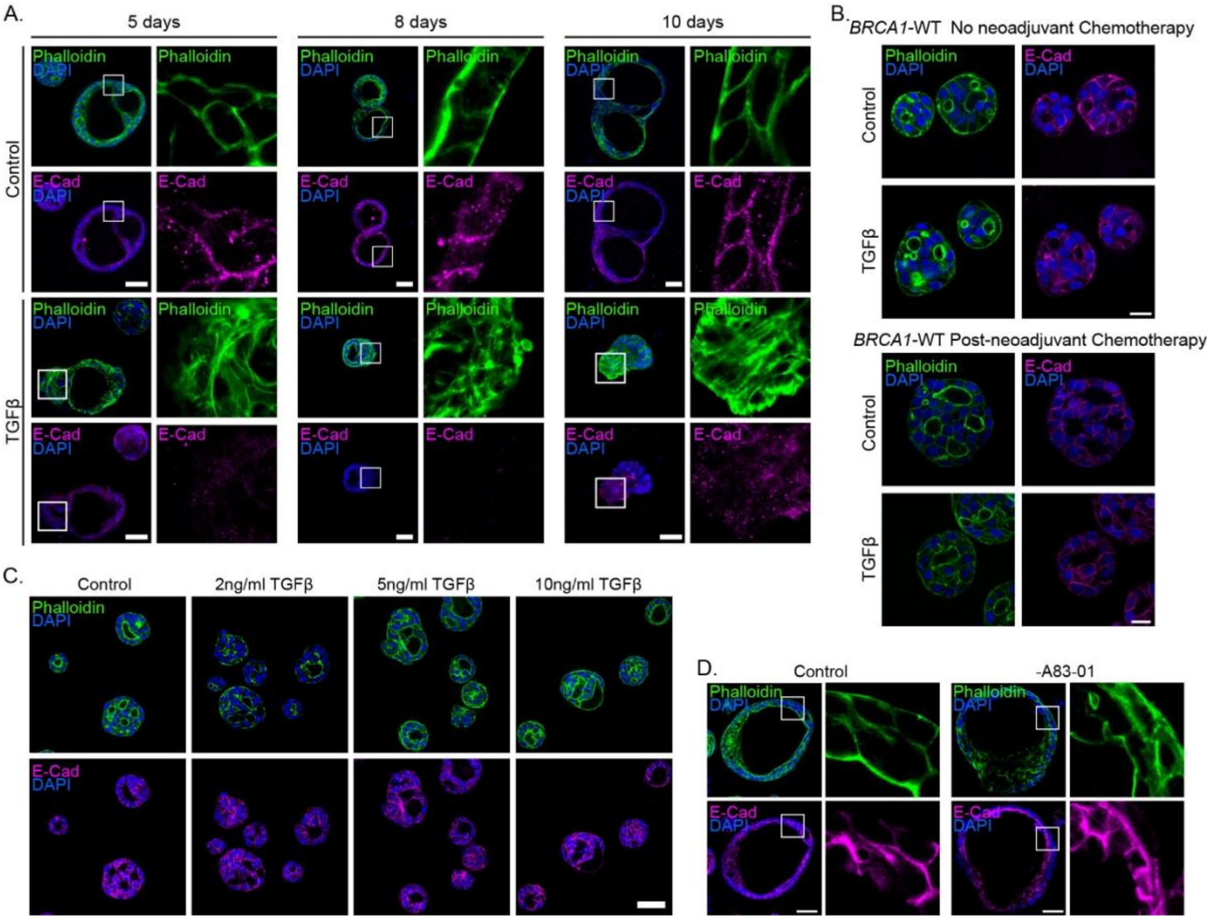


**Supplementary Figure 3. EMT characteristics in *BRCA1*-PV normal mammary gland organoids compared to *BRCA1*-WT.**

A. Time-course analysis of *BRCA1*-PV mammary gland organoids (BR66N). Organoids were treated with TGFβ, and morphological and molecular features were assessed at days 5, 8, and 10. Organoids were immunolabeled for E-cadherin (magenta) and counterstained with phalloidin to visualize F-actin (green). DAPI stain is in blue. Enlargements of the squared areas are shown in right. Top: Control; bottom: TGFβ-treated organoids. Bars=50µm.

B. Confocal images of *BRCA1*-WT mammary organoids derived from patients without (BR21N; upper panel) or with (BR25N; bottom panel) prior exposure to neoadjuvant chemotherapy. Organoids were immunolabeled for E-cadherin (magenta) and counterstained with phalloidin to visualize F-actin (green). DAPI stain is in blue. Top: control; bottom: TGFβ-treated organoids. Bars=20µm.

C. Confocal images of *BRCA1*-WT mammary gland organoids (BR25N) treated with increasing concentrations of TGFβ. Organoids were immunolabeled for E-cadherin (magenta) and counterstained with phalloidin to visualize F-actin (green). DAPI stain is in blue. Bar=50µm.

D. Confocal images of *BRCA1*-PV mammary gland organoids (BR66N) cultured under control conditions or in EMT medium lacking TGFβ (A83-01 withdrawal only; see Methods). Organoids were immunolabeled for E-cadherin (magenta) and counterstained with phalloidin to visualize F-actin (green); nuclei were stained with DAPI (blue). Enlarged views of the boxed regions are shown on the right. Bars=50µm.

### Supplementary Figures

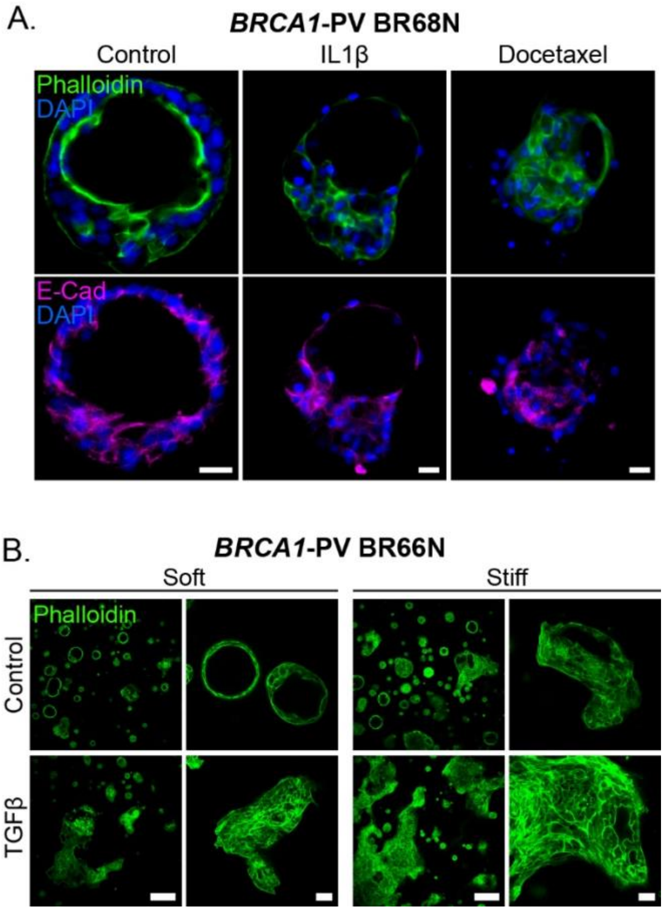


**Supplementary Figure 4. Alternative signals trigger EMT-Phenotype in *BRCA1*-PV organoids.**

A. Confocal images of *BRCA1*-PV (BR68N) normal mammary gland organoids immunolabeled for E-cadherin (magenta) and counterstained with phalloidin to visualize F-actin (green). Organoids were cultured under control conditions or treated with IL1β (30ng/ml) or docetaxel (0.05µM). DAPI stain is in blue. Bars=20µm.

B. *BRCA1*-PV normal mammary gland organoids (BR66N) were cultured in either soft (left panel) or stiff (right panel) matrix and stained with Phalloidin to visualize F-actin. Top: Control organoids. Bottom: Organoids treated with TGFβ. Left: overview images, Bars=250µm. Right: higher-magnification views of individual organoids, Bars=40µm.

### Supplementary Figures

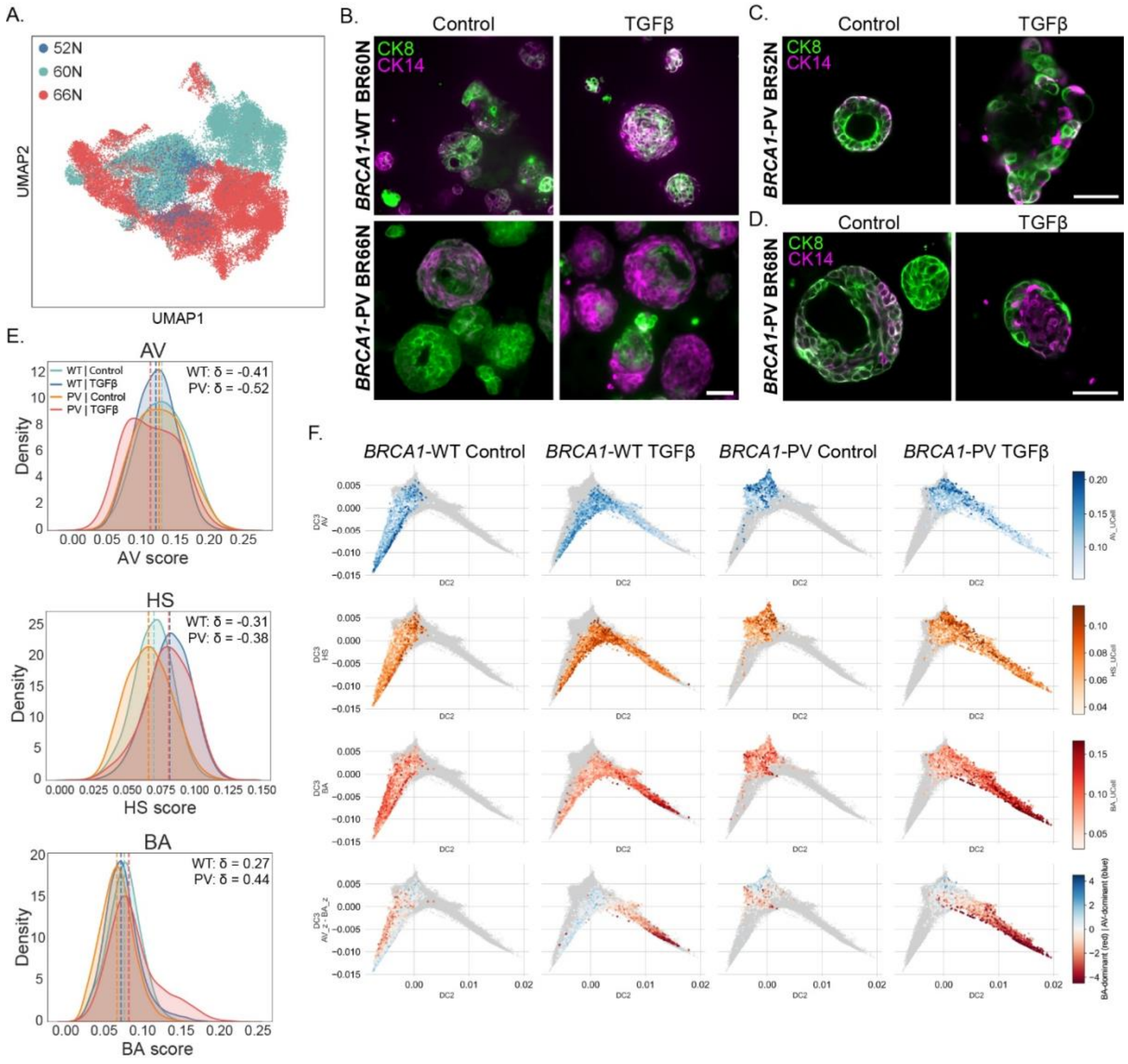


**Supplementary Figure 5. Subtype transitions in *BRCA1*-PV organoids**.
A. UMAP projection colored by sample origin (lines BR52N and BR66N (*BRCA1*-PV); BR60N (*BRCA1*-WT)).

B. FFPE sections of *BRCA1*-WT organoids (top, BR60N) and *BRCA1*-PV organoids (bottom, BR66N) stained by immunofluorescence for CK8 (luminal marker) and CK14 (basal marker) under control or TGFβ conditions. Images demonstrate the organization and expression levels of epithelial subtypes markers within organoid structures. Bar=50µm.

C,D. Confocal images of whole mount *BRCA1*-PV mammary organoid lines (C: BR52N, D: BR68N) stained for CK8 (luminal marker) and CK14 (basal marker) under control or TGFβ conditions. Bars=50µm.

E. Density plots of alveolar (AV), hormone-sensing (HS), and basal (BA) scores; dashed lines indicate medians. Scores were computed using Gray et al. (2022)[43] derived gene signatures. Cliff's δ (Control vs TGFβ within *BRCA1* groups); all $P < 0.001$ (Mann-Whitney, BH-corrected).

F. Diffusion map plots showing the distribution of Alveolar (AV_UCell), Hormone sensing (HS_UCell), and Basal (BA_UCell) scores across *BRCA1*-WT and *BRCA1*-PV organoid groups under control and TGFβ conditions. Cells are displayed on the DC2-DC3 diffusion map embedding, with all cells shown in gray and cells from the indicated group overlaid and colored by UCell score. Bottom row shows a subtype balance index calculated as AV_z - BA_z, where positive values indicate relative AV enrichment and negative values indicate relative BA enrichment.

### Supplementary Figures

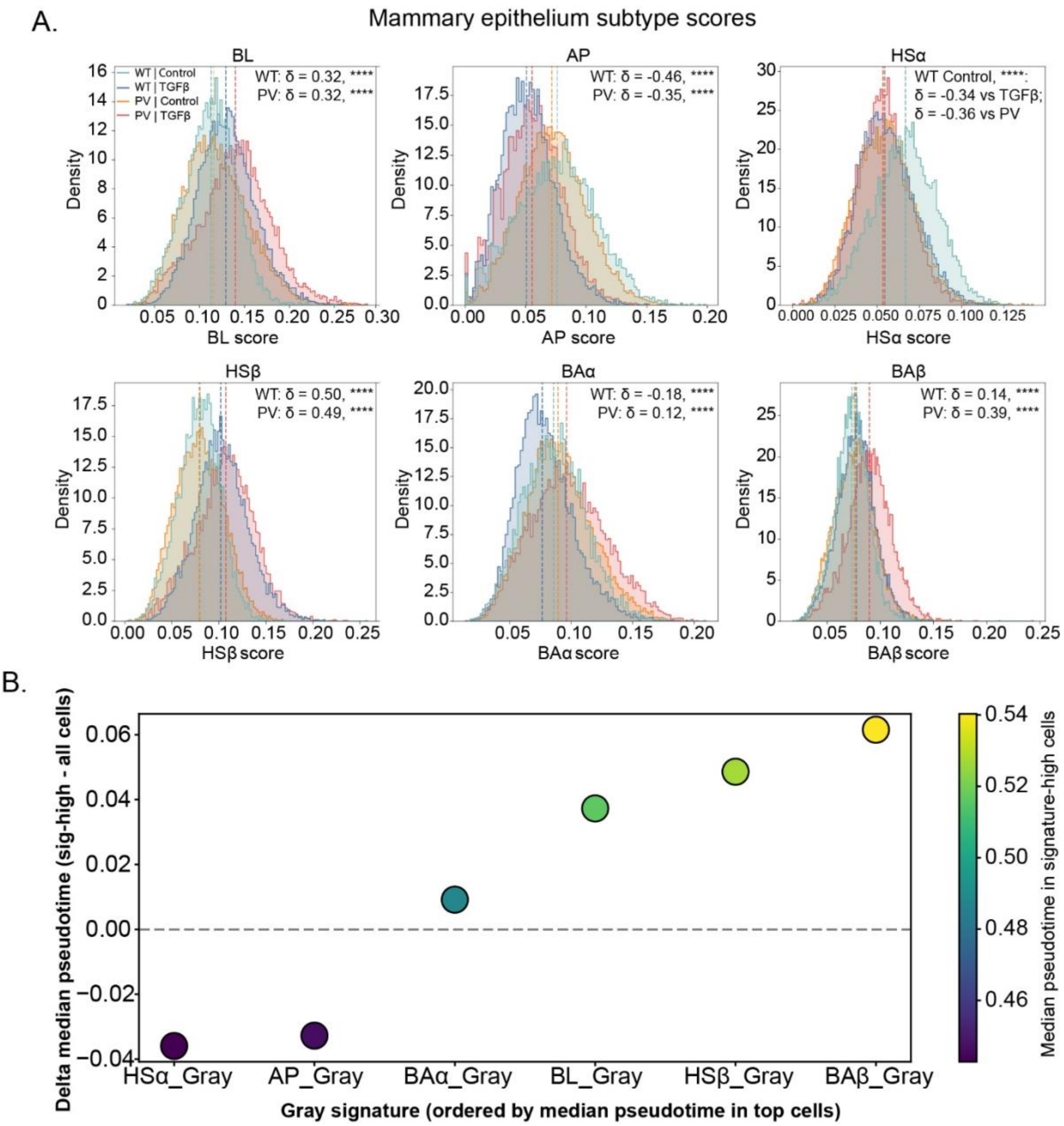


**Supplementary Figure 6. Mammary epithelial subtype signatures shift with genotype and TGFβ exposure and align along pseudotime**.

A. Histograms of single-cell mammary epithelial subtype UCell scores (Gray et al. (2022)[43], grouped by *BRCA1* status (WT, PV) and treatment (control, TGFβ). Distributions are normalized within group; dashed lines indicate medians. Cliff's δ; all $P < 0.001$ (two-sided Mann–Whitney, BH-corrected). BL=basal-luminal, AP=alveolar progenitors, HS=Hormone sensing, BA=Basal.

B. Position of Gray et al. (2022)[43] mammary epithelial subtype signatures along pseudotime. For each signature, the top 20% of cells (by UCell score) were selected. Each point represents one signature, ordered by the median pseudotime of these cells. The y-axis shows the shift in median pseudotime relative to all cells, where positive and negative values denote later and early pseudotime respectively. Color indicates median pseudotime.

### Supplementary Figures

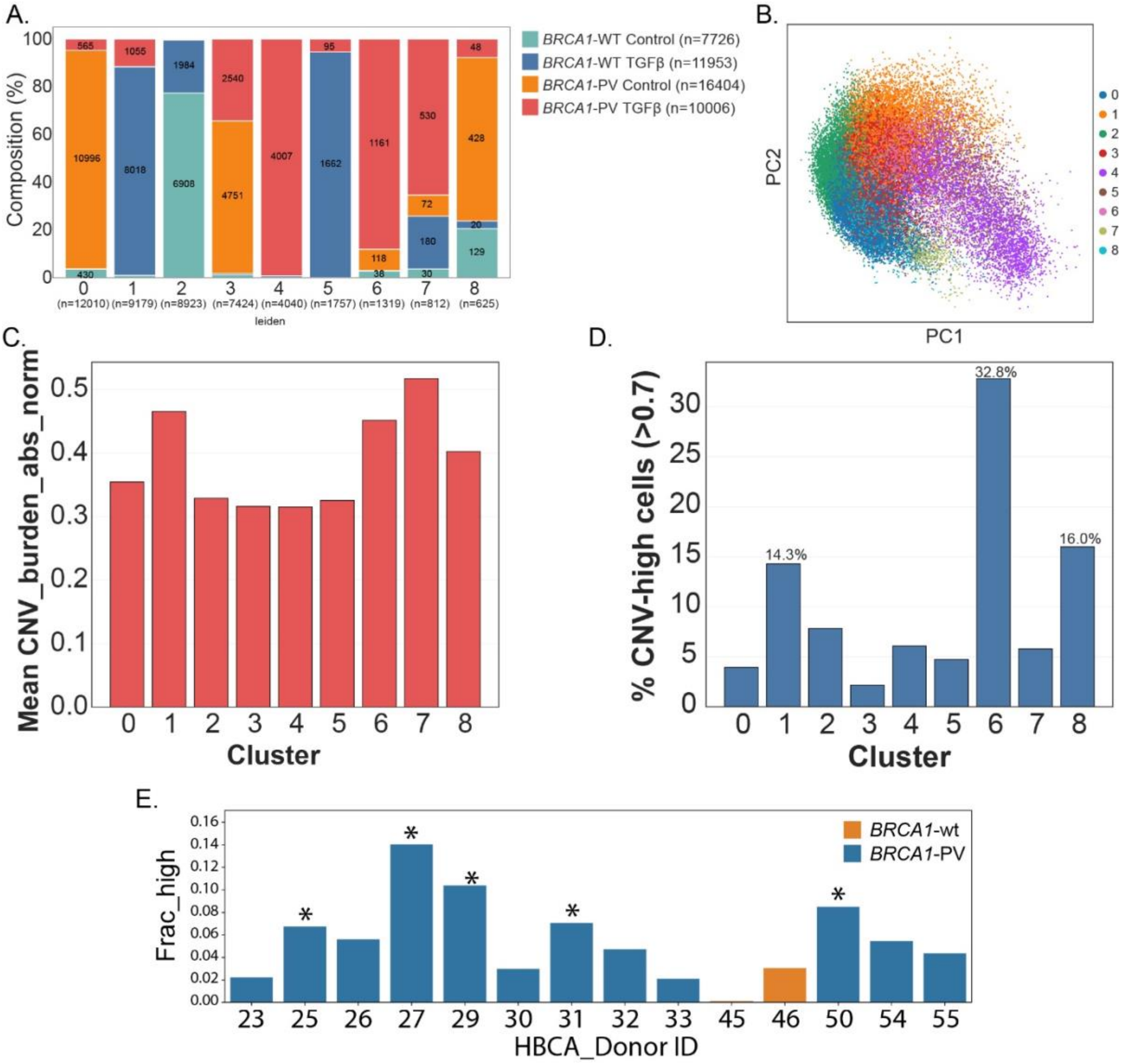


**Supplementary Figure 7. Condition-specific cell states revealed with single cell transcriptomics.**

A. Stacked bar plot showing the proportion of cells from each annotation group (*BRCA1* status and TGFβ treatment condition) within each Leiden cluster. Percentages are shown for segments comprising ≥5% of the cluster. Colors denote annotation groups. n=number of cells.

B. Principal component analysis (PCA) of scRNA-seq data colored by cluster identity.

C. Mean normalized CNV burden per cluster.

D. Fraction of CNV-high cells per cluster, defined using a threshold of ≥ 0.7 on the normalized CNV burden score.

E. Donor-level fraction of cells with high gene signature scores (≥95th percentile) for a *BRCA1*-PV–EMT signature in the Reed et al. (2024)[44] dataset. ~5% are expected to exceed this threshold. Asterisks indicate significant enrichment (binomial test, FDR < 0.05).

### Supplementary Figures

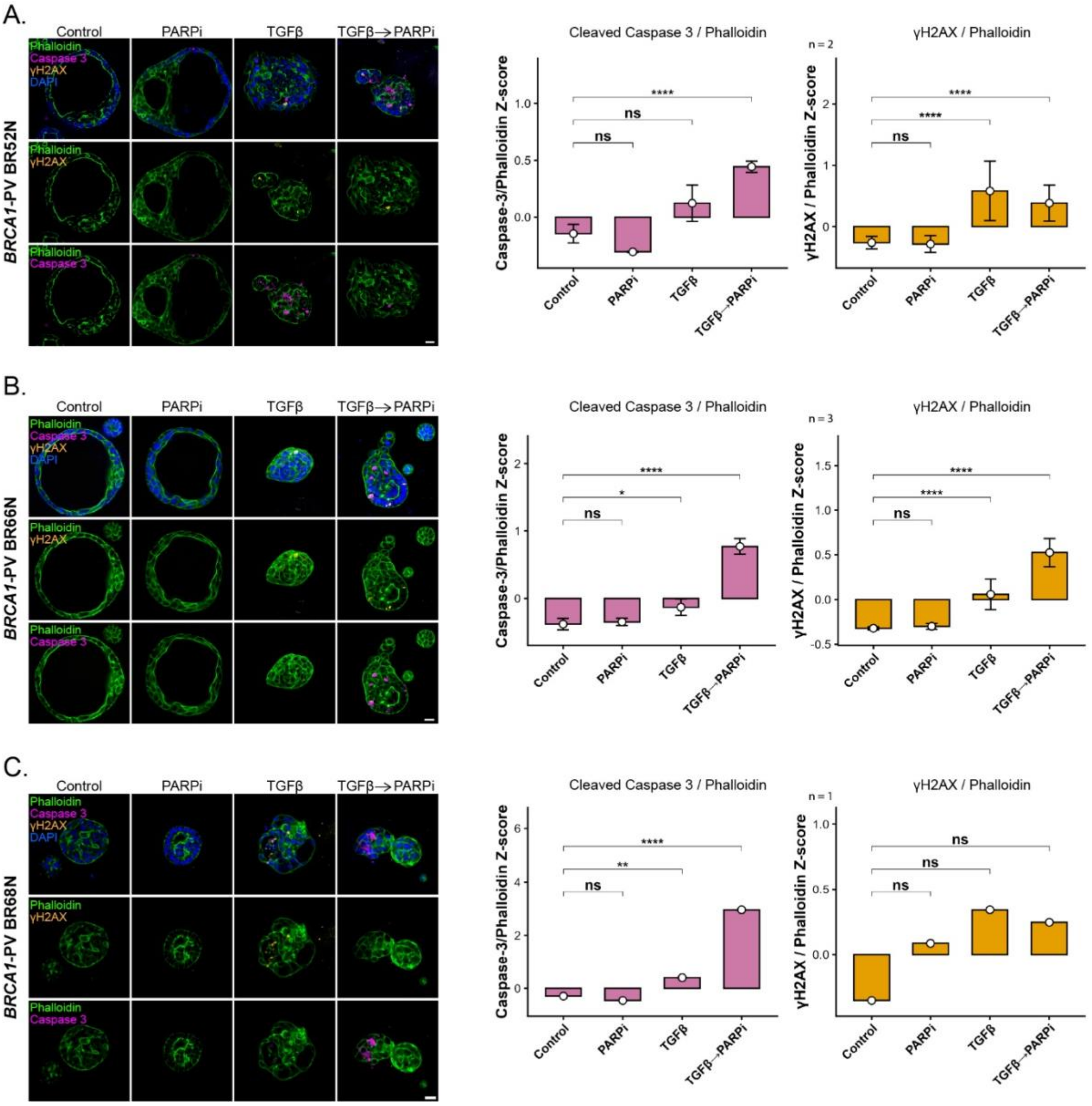


**Supplementary Figure 8. DNA-damage and PARPi susceptibility in *BRCA1*-PV PDOs following TGFβ treatment shown per individual line**.

(A) BR52N, (B) BR66N, and (C) BR68N *BRCA1*-PV organoid lines were treated with DMSO (control), PARPi, TGFβ, or TGFβ → PARPi. Left: Confocal images showing phalloidin (green), γH2AX (orange), and cleaved caspase-3 (magenta) stains. Bars=20μm. Right: Quantification of γH2AX and cleaved caspase-3 intensities measured at the single-organoid level. Marker intensities were normalized to phalloidin to account for organoid size and Z-scored normalized for each biological replicate. Bars represent mean ± SEM across biological replicates within each line. Statistical significance was assessed for each line using linear mixed-effects models with treatment as a fixed effect and biological replicate as a random effect; for BR68N (single biological replicate), linear models were used. Estimated marginal mean contrasts were computed versus control. All lines contribute to the combined *BRCA1*-PV analysis in Fig. 4a. ns, not significant; *$P < 0.05$; **$P < 0.01$; ***$P < 0.001$; ****$P < 0.0001$. n= biological replicates.
